# Interpretable decoding of cell fate from a snapshot of combinatorial signaling

**DOI:** 10.64898/2026.05.17.725652

**Authors:** Abdul-Bassit Fijabi, Seth Teague, Emily Freeburne, Hina Aftab Khan, Craig Johnson, David B. Brückner, Idse Heemskerk

## Abstract

How combinatorial cell signaling controls cellular decisions in the face of crosstalk is a fundamental problem in biology. A key open question is whether a single snapshot of signaling is sufficient to predict cell fate, especially given substantial evidence that signaling dynamics shape fate decisions. Here, we show that a snapshot of combinatorial signaling accurately predicts cell fate at the single-cell level in a model for human embryonic patterning. To this end, we developed Sig2Fate, a quantitative method integrating iterative immunofluorescence, information theory, and machine learning. Cell fate is encoded by combinatorial yet redundant signaling that reduces to a single angular coordinate in the high-dimensional signaling space, providing a simple interpretation of the signal-to-fate map. This map generalizes across variations in BMP concentration and pharmacological perturbations of ERK, Wnt, and YAP signaling, enabling prediction of drug responses from control data alone when signaling crosstalk is accounted for. Our findings provide a framework for predicting and explaining complex phenotypes from signaling perturbations across biological systems.

## Introduction

Cell fate decisions are generally driven by multiple signaling pathways acting in concert, but how cells decode these combined inputs remains poorly understood. This question is central across biology: in embryonic development, multiple morphogens act simultaneously to pattern tissues; in cancer, pathway crosstalk shapes tumor progression and drug response; in immunity, combinations of cytokines control immune response^1–5^. A predictive understanding of these processes requires quantitative maps from combinatorial signaling to cell fate within the spatial context of the tissue, and the ability to predict how cell fates shift when signaling is perturbed pharmacologically or genetically.

Progress has been limited by experimental, conceptual, and computational challenges. Experimentally, measuring many signals simultaneously is difficult: spatial transcriptomics enables highly multiplexed measurements of gene expression^6–8^, but cannot measure cell signaling, which occurs at the protein level through localization and post-translational modification. Meanwhile, spatial proteomics methods are not yet widely accessible. Conceptually, gene expression may depend on combinatorial signals in such a way that no relationship can be discerned by considering one signal at a time^9^. This is illustrated by a toy model in which fate is determined by the difference between two signals such that measuring either signal by itself obscures a simple relationship between signaling and fate (Fig. 1a). Computationally, fate boundaries in high-dimensional signaling space cannot be identified by eye and must be systematically inferred from data. Identifying which signaling combinations are most informative is naturally addressed by information theory but this approach has not yet scaled to large numbers of genes and signals^10–12^.

**Figure 1.**
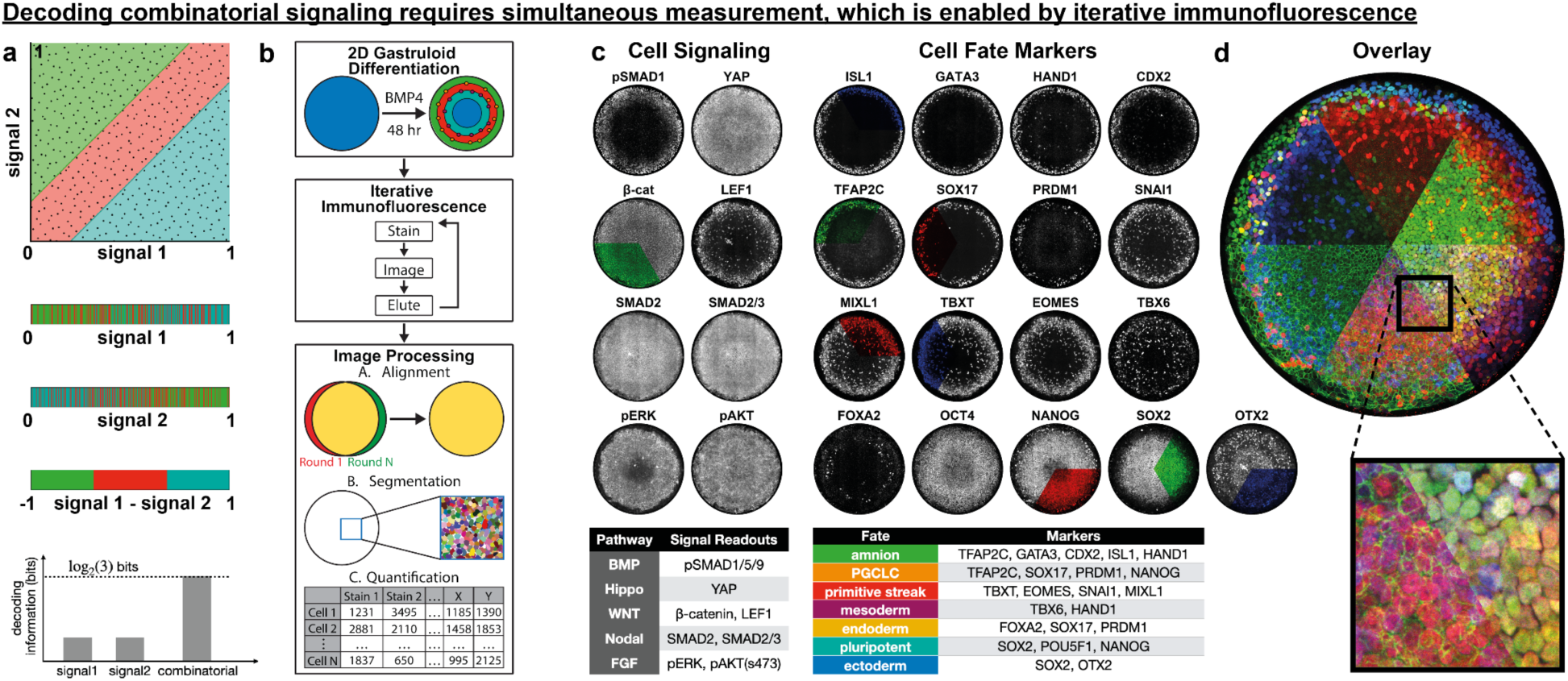
Iterative immunofluorescence enables measurement of combinatorial signaling and cell fate. **(a)** A 2D model where fate depends on the difference in level between two signals illustrates the need to measure combinatorial signaling. Black dots indicating individual cells at the top. Below, 1D projections on signal 1, signal 2 show no clear relationship between signal and fate: the relationship can only be discerned by measuring both signals simultaneously so their difference can be determined (bottom). **(b)** Diagram of our approach. **(c)** Overview of iterative immunofluorescence stains used as proxies for signaling pathway activity and cell fate. Colored sectors of stains indicate color and sector in which these stains are displayed in the overlay in (d). Colors in table below provide an unrelated color code for cell fate followed throughout this manuscript. **(d)** Combination of colored sections from (c). Data can be explored interactively at https://cdb-rshiny.med.umich.edu/2Dgastruloid_IIF/.

Vertebrate gastrulation exemplifies these challenges, as BMP, Wnt, Nodal, and FGF signaling jointly pattern the early embryo. Even for carefully selected direct BMP target genes in zebrafish, the response to BMP changes dramatically when Nodal and FGF are inhibited, illustrating that single-pathway analyses cannot capture how cells decode their signaling environment^13^. Moreover, many studies consider cell population averages, which may mask meaningful (spatial) heterogeneity in a gene-specific manner and cause studies considering different sets of genes to reach different conclusions. As a result, whether cell signaling levels predict cell fate in even the most well-studied patterning processes remains unresolved^13,14^. Due to the inaccessibility of the human embryo, the relationship between signaling and fate is even less clear in human gastrulation. Thus, across systems, fundamental questions remain: What are the combinatorial signaling patterns? Do these explain spatial cell fate patterns at the single-cell level? Can we predict how fate patterns shift when signaling is manipulated?

Here, we address these questions in 2D human gastruloids, an in vitro model for the early human embryo that undergoes self-organized patterning driven by BMP, Wnt, Nodal, FGF, and Hippo/YAP signaling^15–19^. Using iterative immunofluorescence, we collected spatial single-cell data for eight signaling readouts and seventeen fate marker genes, providing a rich reference dataset that can be explored through a public web interface. Combining machine learning and information theory, we then related combinatorial cell signaling to cell fate at the single-cell level. This revealed that a snapshot of combinatorial cell signaling encodes cell fate through a condition-independent, low-dimensional map. The signal-to-fate map predicted cell fate after drug perturbations of ERK, Wnt, and YAP signaling when signaling crosstalk was accounted for. Our findings provide a general framework for relating combinatorial signaling to cell fate and a practical tool for predicting how signaling perturbations reshape cell differentiation.

## Results

### Iterative immunofluorescence enables highly multiplexed measurement of signaling and fate in 2D gastruloids

To relate combinatorial signaling levels to cell type-specific protein expression, we optimized iterative immunofluorescence staining of 2D human gastruloids and developed a computational pipeline to process the resulting image data into a matrix of single-cell protein levels and spatial coordinates^20,21^ (Fig. 1b, Fig.S1ab). We selected a panel of eight antibodies to measure signaling activity, covering the main pathways downstream of the paracrine signals BMP, Wnt, Nodal, and FGF (Fig.1c) as well as the Hippo mechanotransduction pathway. For three of the signaling readouts: YAP, SMAD2, and SMAD2/3, nuclear translocation corresponds to signaling activity so we used the nuclear:cytoplasmic ratio as the quantitative readout, for all others we used the nuclear level. Several of the measured pathway activities are not exclusive to these paracrine signals – for example, ERK is activated by FGF but also by integrin-extracellular matrix signaling – and may therefore convey information about other aspects of a cell’s microenvironment^22^. We further selected a panel of 17 downstream proteins marking all cell types identified in 48h gastruloids^18,23^ (Fig. 1cd) but excluded FOXA2 from subsequent analysis due to negligible expression (Fig. S1e), consistent with our earlier work where endoderm differentiation at 48h was minimal^18^. A prerequisite for relating signaling to fate is that measurement noise is small relative to biological variation. We confirmed this by a repeated stain for SOX17 in rounds one and seven, which yielded R^2^=0.97 implying only 3% of the overall variance in protein levels is due to technical noise (Fig.S1cd). Thus, we obtained a rich, high quality dataset, which can be explored through an online visualization tool (see data availability). These data enabled us to address several open questions: Can cell fate be predicted from signaling at the single-cell level? Which signals or combinations carry information about each fate marker, and does the signal-to-fate map have a simple, interpretable structure?

### Combinatorial signaling predicts gene expression patterns

To determine whether combinatorial signaling predicts cell fate, we trained a machine learning model to predict fate marker gene expression from signaling measurements alone. Our signal-to-fate model, Sig2Fate, uses a variational encoder-decoder network with a bottleneck that compresses the map into a low-dimensional latent space (Fig. 2a, Methods). Sig2Fate was trained on six colonies using leave-one-out cross-validation at the colony level. We optimized hyperparameters and controlled for overfitting (Fig.S2), after which Sig2Fate substantially outperformed linear regression and matched the performance of random forest regression, the best other non-linear alternative (Fig.S2a-c). Predicted gene expression patterns agreed with experimental measurements in both location and magnitude (Fig. 2b-f), including strong agreement in the conditional average <g_exp_ | g_pred_> across all genes (Fig. 2g, Fig. S3a-f). Furthermore, we found that the model captures the key correlations between different genes in single cells, including correlated expression of genes marking a single fate, and anti-correlated expression of genes exclusive to distinct fates (Fig. 2h).

**Figure 2:**
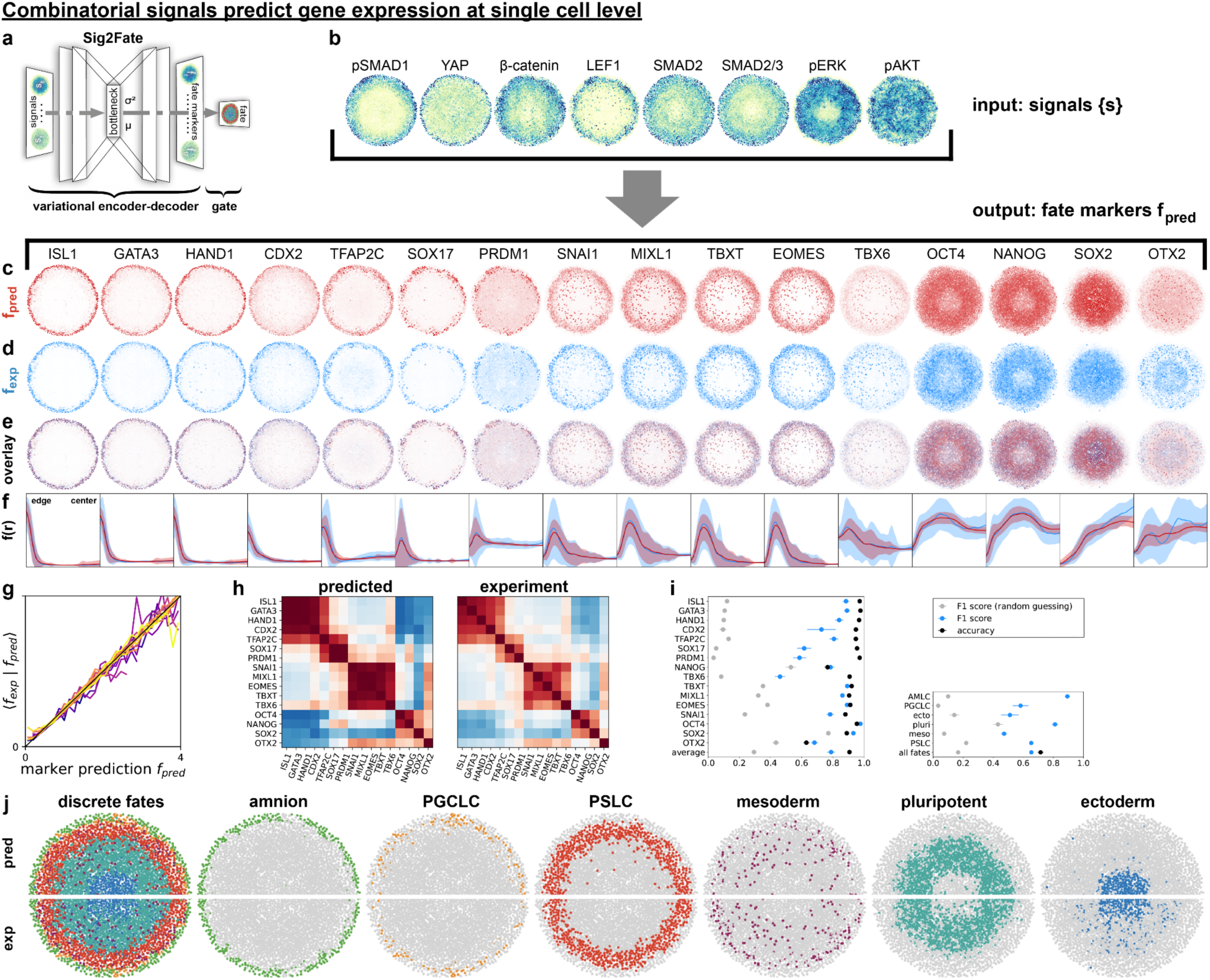
Combinatorial signaling activities predict spatial patterns of gene expression. **(a)** Machine learning architecture used to predict fate from signaling. **(b)** Eight single-cell signaling activities in the same colony, which serve as input for the signal-to-fate prediction model, which was trained with 6-fold cross-validation where each colony is predicted from the others. **(c)** Predicted expression for 16 marker genes in the same colony. **(d)** Measured expression in the same colony. **(e)** Overlay of predicted (red) and measured (blue) positive cells. **(f)** Predicted (red) and measured (blue) mean radial gene expression profiles. Shaded regions correspond to standard deviation of single-cell gene expression around the mean. **(g)** Conditional average of the measured gene expression as a function of predicted gene expression for all marker genes (colors). **(h)** Predicted (left) and measured (right) Pearson correlation coefficient between pairs of marker genes. **(i)** F1 score and accuracy of discretized (positive or negative) prediction of each marker gene (left) and for fate assignment (right). Error bars: standard deviation over N=6 colonies. **(j)** Predicted (top) and measured (bottom) cell fate for a representative colony, mirrored in the midline. PGCLC: primordial germ cell-like cell; PSLC: primitive streak-like cell.

To evaluate performance at the single-cell level, we classified cells into positive and negative for expression of each gene, and found an average overall accuracy of 0.9 (90% correct prediction) (Fig. 2i, Fig.S3g, Methods). A more stringent metric is the F1 score (Methods), which balances how many positive predictions were correct (precision) with how many of the actual positive cells were predicted (recall). F1 scores exceeded 0.8 for most genes, including all amnion- and primitive-streak-associated markers. Scores were lower for scattered cell types: approximately 0.6 for SOX17+ and PRDM1+ primordial germ cell-like cells (PGCLCs) and 0.45 for TBX6+ mesoderm, though still much higher than expected from a random draw (Fig.2i). By combining these thresholded gene-wise predictions (Methods), we classified cells into discrete fates with overall accuracy and F1 score of 0.72 and 0.65, respectively. Performance was higher for pluripotent cells, primitive streak (PS)-like cells, and amnion-like cells but lower for mesoderm, PGCLCs, and ectoderm. Model performance was similar between experimental repeats (Fig.S3h) and captured the correct spatial distribution of all 6 cell fates (Methods; Fig. 2j). In summary, cell fate can be predicted from combinatorial signaling to a remarkable degree.

### Cell fate is encoded combinatorially but redundantly across signaling pathways

The efficacy of the signal-to-fate map raises the question: which signals are required to predict expression of each fate marker? Is each gene predicted by a single signal, or are multiple signals required in combination? We quantified this with mutual information (MI) between signals and fate, I({s}, f), which measures how well signals predict fate in bits and, unlike correlation, captures nonlinear relationships^24^. Fig. 1a illustrates this for the toy example: the MI between each individual signal and fate is low, while the MI of their combination equals log₂(3) bits as expected.

We first calculated the pairwise MI of signals and fate markers (Fig. 3a). This revealed a complex network of relationships: some genes such as ISL1 and TBXT exhibit dominant contributions from single signals (pSMAD1 and LEF1 respectively), while others such as NANOG and SOX2 exhibit similar MI with many signals (Fig. 3a, Fig.S4ab). However, pairwise MI does not show whether different signals provide independent or redundant information, nor does it reveal synergistic information like in the toy model in Fig. 1a. Pairwise signal-signal MI provides a first hint at redundancy: MI was highest between SMAD2 and SMAD2/3 and significant between pSMAD1 and readouts of the downstream Nodal and Wnt pathways, suggesting substantial redundancy in the information they contain about fate marker expression.

**Figure 3:**
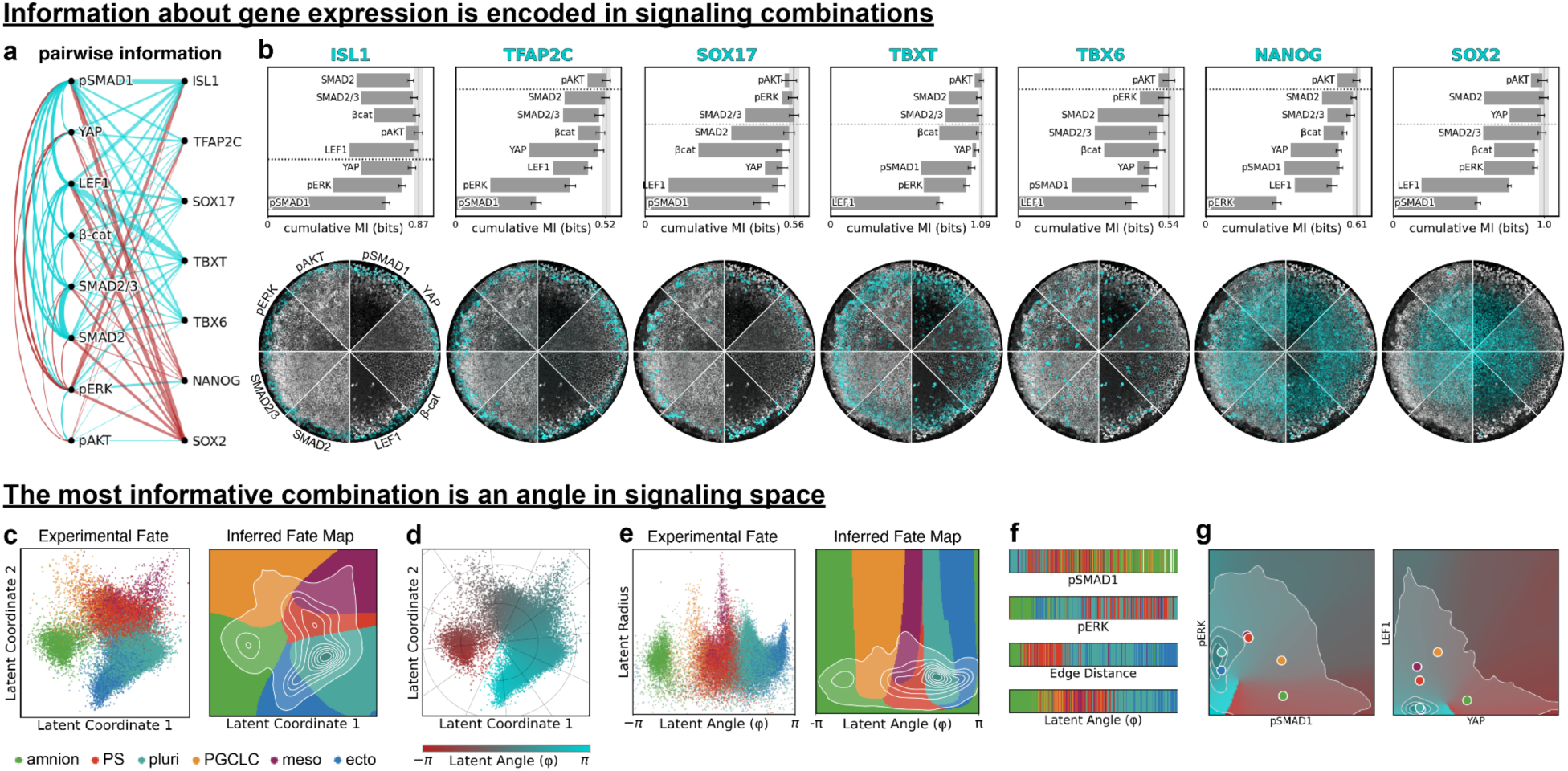
Combinatorial signaling encodes cell fate through a single angular coordinate. **a)** Graph of signal-fate relationships, line weight represents mutual information, color indicates positive (cyan) or negative (red) correlation. **b)** Top row: cumulative mutual information between signals and fate markers visualizes redundancy. Ordered from most informative to least informative signals, above the dashed lines the total information is no longer significantly increasing relative to the maximum. Bottom row: overlay of signaling stains (pie slices) with each marker genes for representative colony. **c)** Data in latent coordinates colored for cell fate. Left: scatter plot of full data set colored by experimentally classified cell fate (N=6 colonies, n=30431 cells). Right: Colormap of predicted fate at each location in latent space, contours show probability distribution of full data set. **d)** Data in latent coordinates overlaid with polar coordinate grid and colored for polar angle. **e)** Data in latent polar coordinates colored for cell fate, colors as in (c). **f)** Bar plot showing fate for randomly sampled cells in equally spaced bins across the range of each signal, N=450 cells total, upper and lower limits are 1st and 99th percentiles. Fate colors as in (c). **g)** Heatmap of latent angle for two pairs of input signals evaluated at the conditional mean for all other signals, overlaid with kernel density estimate of data density (contour lines) and the mean of each fate (colored dots, colors as in (c)).

To determine how much unique and redundant information each signal contains about each fate marker, we sought to calculate MI between combinations of signals and gene expression. However, accurate MI estimation requires well-sampled joint probability distributions, which becomes prohibitive in high dimensions^25,26^. Machine learning provides a solution: because the model acts as a decoder of signals into fate, the MI between prediction f* and measurement f provides a lower bound, I({s}, f) ≥ I(f*, f), by the data processing inequality^24,27^. Consensus across model architectures suggests this bound is tight^27,28^. We therefore leveraged Sig2Fate to estimate MI between cumulative combinations of signals and fate: starting with the most informative single signal for each gene, we sequentially added the signal that increased the total MI the most, testing at each step whether the increase was significant (Fig.3b, Methods). This revealed that combinatorial signaling is essential to explain fate marker expression: for most genes, more than two signals were required to reach the total MI and which combination of signals was most important differed between genes (Fig. 3b, Fig. S4cd). Combinatorial signaling was most important for TFAP2C and NANOG: the most informative signal contributed only about 50% of the total information and all signals except pAKT contributed significantly to the total. To demonstrate the implications of this combinatorial decoding, we predicted gene expression from pSMAD1 alone, which has been claimed to explain the entire fate pattern^29^. This performed much worse by all measures and missed two out of six cell fates (Fig. S5).

Although a signaling snapshot need not reflect causal relationships between signals and fate markers, the MI analysis aligned strikingly with known gene regulation mechanisms. For example, amnion differentiation is primarily controlled by BMP signaling and does not require Wnt or Nodal^28,30^. In line with this, pSMAD1 contained 78% of the information about the amnion maker ISL1 and no significant contributions were made by Wnt and Nodal readouts (Fig.3b). Similarly, TBXT which is a direct target of Wnt signaling^31^ and accordingly most information (72%) about this gene comes from the Wnt readout LEF1, with the next biggest contribution from pERK (to 92%), which is also required for TBXT expression^32–34^. Surprisingly, although Nodal strongly regulates TBXT, Nodal signaling readouts do not add information to LEF1 and pERK, suggesting the information carried by Nodal is fully redundant with these signals in encoding TBXT. Note that high MI is also expected when a signal anticorrelates with a fate marker, which explains pERK as the second largest contributions for ISL1: ERK signaling is lowest in the amnion. We used these insights to determine a reduced set of signals for predicting cell fate, simplifying follow-up experiments by removing the need to stain for all signals. pAKT contained very little information about fate overall and made no significant contribution for most genes (Fig. 3b, Fig. S4a-d). Furthermore, contributions of SMAD2 were highly redundant with SMAD23 (Fig. 3b, Fig. S4a-d). Thus, removing pAKT and SMAD2 had only a small effect on model performance (Fig. S4e) and we subsequently used a reduced model with six signaling readouts.

Although different signals provide complementary information, there is also significant redundancy: the sum of individual signal-fate marker MIs greatly exceeds the MI of all signals combined (e.g., 2.35 vs. 0.97 bits for SOX2; Fig. S4f,g). Together, these results demonstrate that fate is encoded in inherently combinatorial, yet highly redundant cell signaling.

### Cell fate is encoded by an angle in signaling space

The MI between signals and fate depends on both the signal-fate relationship and the distribution of the data in the signaling space. However, Sig2Fate learns the underlying relationships in the form of a condition-independent signal-to-fate map that can be interpreted by relating its bottleneck coordinates to the measured signals. To validate this approach, we returned to the toy model of Fig.1a, where fate depends on the difference between two signals (Fig.S6a). Although different data distributions strongly affected the MI between signals and fate markers (Fig.S6b,c), Sig2Fate mapped data from all conditions onto a single line in latent space, and polynomial approximation confirmed that the latent coordinate along this line is proportional to the signal difference (Fig.S6d,e). Thus, Sig2Fate correctly recovered the underlying signal-to-fate map from data across conditions.

We then similarly analyzed the signaling code of 2D gastruloids. Although the signaling data occupied the full 6D signaling space, the signal-to-fate map could be compressed into a 2D latent space without loss of predictive power (Fig. S2b, Fig.S7a). This implies that many distinct signaling combinations map to the same fate marker expression. The data formed an arc along which cell fates were ordered similar to their spatial arrangement from colony edge to center, while inferred fate boundaries carved out wedges of the space (Fig. 3c). This motivated us to introduce polar coordinates (φ,r), where the angle φ traverses the arc (Fig.3de). Analogous to the signaling difference in the toy model (Fig.1a), φ separated fates better than individual signals or edge distance: for example, amnion and primordial germ cells overlapped in edge distance, but were well-separated by φ, and mesoderm had a broad spatial distribution but occupied a narrow range of φ (Fig.3f, Fig.S7bc). Fate prediction based on φ alone performed only slightly worse than prediction using the full latent space, although r was required to separate mesoderm from primitive streak (Fig.3e, Fig.S7d); consistently, reducing the latent space to 1D yielded a coordinate that approximated φ (Fig.S7e). Signaling profiles across the latent space recapitulated known activity: for example, pSMAD1 was highest in the amnion and pERK in the primitive streak (Fig.S7fg).

Visualizing the latent coordinate against pairs of signals revealed the arc-like structure of the latent space was also present in the input signal space, rotated and deformed depending on the projection plane (Fig.3g, Fig. S7h). This suggested that the latent angle corresponds to an angle in high-dimensional signaling space. Indeed, we identified an angle in the full signaling space that corresponds directly to the latent angle (Fig.S7ij, Methods). In contrast, we found no simple (linear) relationship between the latent radius and the true radius in signaling space (Fig.S7k).

In summary, whereas in our toy model there was a simple linear relationship between fate and signaling, the relationship between signals and fate in the 2D gastruloid is a function of an angle in the signaling space, which is a highly non-linear function of the signals itself. Consistent with the combinatorial nature of the code identified by our MI analysis, the MI between cell fate and latent angle is much stronger than with any individual signal. Taken together, these results demonstrate that signals map to cell fate through a combinatorial and redundant, yet interpretable and low-dimensional mapping.

### The signal-to-fate map is conserved when signaling changes

The signal-to-fate map is only biologically meaningful if it is conserved across different signaling conditions. In other words, when signaling is changed, does cell fate change in a consistent and predictable manner?

To address this question, we induced altered signaling patterns by changing the exogenous BMP concentration away from the standard value of 50 ng/ml (‘B50’). Specifically, lowering the BMP concentration to 10 ng/ml (‘B10’) led to a shorter-range pSmad1 gradient, while a concentration of 200 ng/ml (‘B200’) caused BMP signaling to extend further into the colony (Fig. 4a). The altered BMP signaling caused all downstream signals to also change (Fig. 4a). For instance, B10 exhibited a more pronounced Smad2/3 gradient, while B200 had the opposite effect and led to almost uniform Smad2/3 activity.

**Figure 4:**
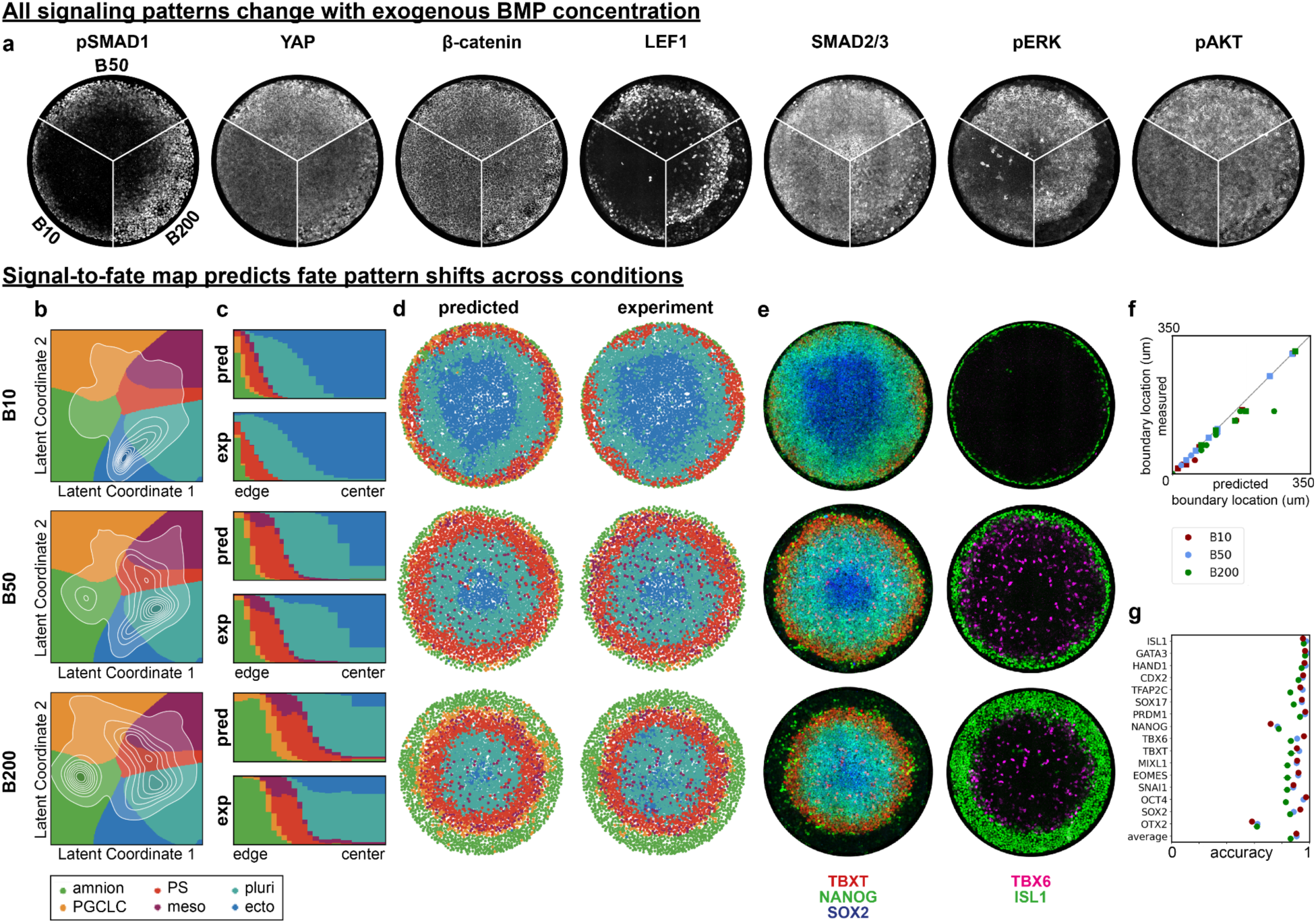
The signal-to-fate map is conserved when signaling changes. **(a)** IIF images of all 7 signals for three different exogenous BMP4 concentrations. **(b)** Predicted latent space distributions for different BMP concentrations. Predictions are performed using the same model trained only on B50 data that is also used in Figs. 2,3. **(c)** Predicted (pred) and measured (exp) spatial distribution of cell fates as a function of radius from edge to center. **(d)** Predicted (left) and measured (right) discretized cell fate patterns for different BMP4 concentrations in representative colonies. **(e)** IIF stains of fate marker expression patterns for the same colonies as in (d) in each condition. **(f)** Predicted versus measured fate boundary locations. **(g)** Prediction accuracy of discretized gene expression for all three conditions. Statistics are for N=6 colonies per condition.

When we applied our model trained on B50 to the measured signals from B10 and B200 conditions, it predicted large shifts in the relative proportions and spatial extent of the cell fate domains (Fig. 4b-d): in B10, a much larger central ectoderm region surrounded by a broad ring of pluripotent cells was predicted, while all other cell fates exhibited smaller domains closer to the edge. In B200, in contrast, the model predicted a broad amnion region enclosing a smaller primitive streak ring and a central pluripotent domain without significant ectoderm. Remarkably, these predictions quantitatively captured the fate domain shifts observed experimentally (Fig. 4de, Fig.S8a). Even at the single-cell level, the prediction accuracy of individual genes in B10 and B200 was comparable to the B50 prediction (Fig. 4f), and did not decrease much relative to models trained on each individual condition, although F1 scores were expectedly poor for markers and fates that were (nearly) absent in the perturbed conditions, like TBX6 and mesoderm for B10 (Fig. S8b-f). Thus, Sig2Fate generalizes across conditions, suggesting a conserved signal-to-fate map.

### Signal-to-fate map predicts the outcome of signaling pathway perturbations

Given that our model generalized to different BMP concentrations, we asked whether it could also predict the effect of pharmacological perturbations of specific pathways. We tested the B50 signal-to-fate map against three drugs targeting distinct pathways: WntSeci (IWP2, Wnt secretion inhibitor), a MEKi (MEK inhibitor, blocking ERK phosphorylation), and LATSi (TRULI, LATS inhibitor, activating YAP) (Fig.S9a-c). When given measured post-perturbation signaling profiles (Fig.5ab), the model correctly predicted fate for MEKi, and largely WntSeci, confirming that the signal-to-fate map is conserved across these perturbations (Fig. 5c-e, S9d). Interestingly, both of these perturbations mapped across the previously unsampled region between the ends of the arc-shaped distribution in latent space, demonstrating that this region is biologically meaningful (Fig. 5f).

**Figure 5:**
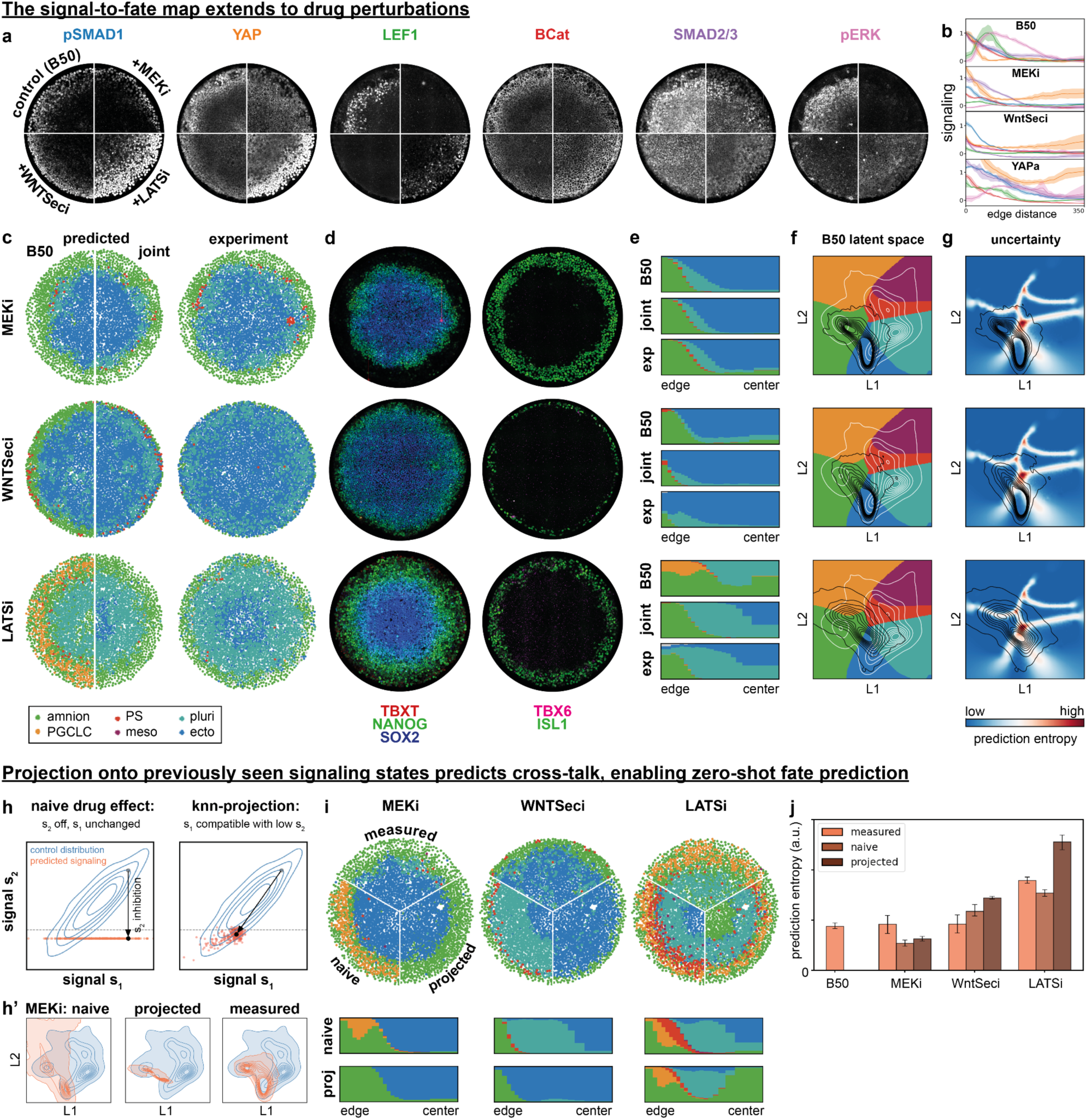
The signal-to-fate map predicts the fate outcome of signaling perturbations. **a)** Stains for signaling activity after drug treatment compared to control. **b)** Radial signaling profiles corresponding to (a). **c)** Fate predictions from measured signaling activity after drug treatment with either the same B50 model as in Fig.2-4, or a combined model trained on B50 and the drug treatment (joint). **d)** IIF stains of fate marker expression patterns for the same colonies as in (c). **e)** Spatial distribution of cell fates as a function of radius from edge to center as predicted from the B50 or joint models, or measured (exp). **f)** Overlay of control and treated distributions in the B50 latent space shows LATSi peaks close to fate boundaries. **g)** Overlay of prediction uncertainty (entropy) and distributions of drug perturbed conditions show uncertainty peaking where boundaries meet. **h)** Cartoon illustration of k-nearest neighbor (knn) projection to account for signaling crosstalk. **h’)** Effect of knn-projection on the latent space distribution for predicting the effect of MEK inhibition (setting ERK low). **i)** Naive and projected predictions of cell fate for each drug perturbation compared to prediction from measured signaling. **j)** Prediction entropy indicates LATSi predictions are unreliable. N=6 colonies for all conditions except LATSi, which is N = 4.

For LATSi, however, the model failed (Fig. 5c-e). To understand why LATSi was not predicted by the B50 map, we quantified prediction uncertainty across the latent space (Fig. 5g). For each cell, we sampled repeatedly from its learned latent distribution, which reflects how precisely the model could place it on the fate map, and computed the entropy of the resulting fate predictions. This gives a per-cell measure of how robustly the model assigns a fate: low entropy means the same fate is predicted every time, high entropy means the model is uncertain. As expected, uncertainty was higher near fate boundaries and peaked near the origin of the polar coordinates where fate boundaries converge. LATSi cells land disproportionately in this region (Fig.5g), explaining why the B50 model produces unreliable predictions for this condition. Training a joint model on B50 and LATSi data restored prediction accuracy for both conditions, demonstrating that a single conserved map exists (Fig. 5c-e). Importantly, this joint model does not simply refine the fate boundaries near the origin: instead, it defines a new latent space where LATSi cells are shifted relative to, and partially outside, the B50 fate domains (Fig. S9e), indicating that LATSi activates a signaling regime not represented in B50 alone.

Having established when the map generalizes, we asked whether fate outcomes could be predicted without measuring the perturbed signaling profiles. Such a zero-shot prediction would demonstrate that Sig2Fate is practically useful for predicting drug perturbations. Naively setting each drug’s primary target to an extreme value while holding other signals at B50 levels failed for all four drugs (Fig.5i), underscoring the pervasive role of signaling crosstalk (Fig. 5ab). We therefore sought to predict the perturbed signals including crosstalk. We hypothesized that the natural heterogeneity in B50 data encodes the biologically compatible signaling states. For each cell, we identified its nearest neighbor in B50 signaling space whose target signal was already in the drug-induced state: below a threshold representing pathway suppression for inhibitors, or above a YAP threshold for LATSi. We then substituted that cell’s signaling profile as input to the model (Fig.5h). For MEKi and WntSeci we excluded pSMAD1 from this procedure, reasoning that as a response to an exogenous signal that is upstream of these signals it is subject to minimal crosstalk. As an example, this procedure captured the known relationship between Nodal and Wnt signaling: if LEF1 is low, SMAD2/3 will be low because Nodal is downstream of Wnt. Indeed, for WntSeci predictions were much better and for MEKi, the zero-shot prediction matched experimental fate patterns nearly perfectly. For LATSi the projection failed, consistent with LATSi inducing signaling combinations not present in B50 (Fig. 5i). Prediction entropy of the projected signaling aligned with that of measured signaling, with LATSi showing the highest uncertainty and MEKi the lowest (Fig. 5j). Thus, prediction entropy can be used to determine when a perturbation moves the system into a region where the model lacks support and additional data is needed.

Together, these results establish a general framework for extending Sig2Fate to new pharmacological contexts: natural signaling heterogeneity in control data enables zero-shot fate prediction through crosstalk approximation, while prediction entropy identifies the conditions where this strategy breaks down, prioritizing them for targeted experimental follow-up.

## Discussion

Here, we showed that a snapshot of combinatorial cell signaling encodes cell fate through a conserved, low-dimensional signal-to-fate map in 2D human gastruloids. Cell fate is encoded in an inherently combinatorial yet highly redundant signaling state: most fate markers required more than two signals for maximal prediction, with different combinations relevant for different markers, yet the information carried by different pathways overlapped substantially. The compression of the signal-to-fate map into a low-dimensional latent space reflects both this redundancy and the existence of signaling directions that carry little information about fate (like in the toy model in Fig.1a). Cell fates are arranged in signaling space along an angular coordinate φ that carries more fate information than any individual signal. This can be viewed as a multi-signal generalization of the classical gradient-threshold model of positional information^35,36^ where here, thresholds depend on a coordinate constructed from the combined activity of multiple pathways. The angle is also recovered in the signaling space itself and optimally predicts cell fate, indicating that it reflects the underlying geometry of the signal-to-fate map rather than an artifact of dimensionality reduction^37^.

Similar low-dimensional motifs have been identified in diverse systems driven by combinatorial inputs, from early fly patterning^38^ to immune cell responses^4^. In mammary epithelial cells, combinatorial signaling responses after brief growth factor treatment have been shown to encode cellular state heterogeneity^39^. The encoding we observe occurs in a different regime: developmental patterning after multiple days of differentiation, where combinatorial signaling controls spatial patterning on a tissue scale. In this setting, the redundancy between pathways and compression to a low-dimensional space may provide robustness to noise and to perturbation in individual pathways.

A practical benefit of the conserved signal-to-fate map is that it can be used to predict the outcome of perturbations the model has never seen. Using measured post-perturbation signaling as input, the model correctly predicted fate outcomes for altered BMP concentrations and inhibition of Wnt and ERK signaling, confirming that the map is conserved across these perturbations. YAP activation was the only exception, but incorporating it into the training data restored prediction accuracy across all conditions, demonstrating that the map itself remains consistent — the control condition simply did not sample the signaling states that pharmacological YAP activation produces. This has an interesting biological interpretation: whereas BMP, Wnt, FGF, and Nodal form a well-characterized signaling hierarchy driving gastrulation, the role of Hippo/YAP signaling within this hierarchy remains unclear. The fact that LATSi drives cells into signaling states outside the natural variation of the unperturbed system suggests that YAP activation engages a qualitatively distinct signaling network, indicating that Hippo/YAP signaling sits outside the established gastrulation hierarchy rather than acting as one of its components.

That a model trained on control cells generalizes to pharmacological perturbations implies that zero-shot prediction is possible if the perturbed signaling profile can be estimated. The central challenge is signaling crosstalk: perturbation of one pathway reshapes the activity of others, so that naive predictions assuming a drug affects only its direct target fail. However, successful prediction from measured post-perturbation signaling implies that the resulting signaling states are already present in the control population, so that perturbed signaling can be approximated without measurement by finding control cells whose signaling matches the expected post-perturbation state. This was the case for ERK and Wnt inhibition, where zero-shot predictions based on control data alone closely matched experimental fate patterns. Furthermore, prediction entropy provides an intrinsic confidence metric that flags conditions where predictions are unreliable, so these can selectively be added to the training data to restore accuracy. Together, these results define an iterative strategy: control data enables initial fate predictions, prediction entropy prioritizes conditions requiring experimental follow-up, and targeted measurements extend the map to new regions of signaling space. This approach extends naturally to any system where combinatorial signaling drives a phenotypic transition, and differs from methods that train directly on drug-phenotype pairs by instead learning the underlying signal-to-fate map^40^.

A snapshot-based encoding may seem to conflict with prior work showing that signaling history, not instantaneous level, determines fate^28,41–47^. We propose that although fate is determined by the temporal history of signaling, a combinatorial snapshot can contain equivalent information about fate. For example, the history of BMP signaling is required to explain amnion differentiation from BMP alone, but a snapshot of combinatorial signaling, including elevated YAP and reduced ERK in amnion relative to pluripotent cells, may encode the same information^28,48^. This relationship is in principle correlative and could partly reflect fate influencing signaling rather than the reverse. Yet our model correctly predicts that ERK inhibition and YAP activation promote amnion differentiation, demonstrating that these signals are causally upstream of fate. This raises an interesting possibility: in early gastrulation, different cell types may actively reinforce the signaling that specifies them through positive feedback. If these signals act non-cell-autonomously, this would imply a community effect in which cells make spatially correlated fate decisions^49,50^. Establishing the structure of these feedback loops will require dynamic perturbation experiments.

An essential question for future work is how the signal-to-fate map evolves over time: as later fates emerge, is the map extended or does the mapping itself change? Our framework also raises theoretical questions about whether the observed crosstalk relationships are structured to maximize combinatorial information and what properties make signal-to-fate maps robust to variation in input signals, questions that apply not only to gastrulation but to any system where multiple signals jointly specify cell fate. Our work provides a general strategy for dissecting how combinatorial signals control complex phenotypic transitions, with potential applications ranging from cancer, where pathway crosstalk confounds targeted therapy^5,51,52^, to synthetic morphogenesis, where engineering predictable fate patterns requires quantitative maps from signals to outcomes^53^.

## Acknowledgements

We thank Gašper Tkačik and Alex Schier for inspiring discussions. This work was supported by NSF RECODE (2033654, IH) and the National Institute of General Medical Sciences (NIGMS R35GM138346, IH). BF was supported by the Michigan Postbaccalaureate Research Education Program (NIH R25 GM086262). EF was partially supported by the NIH Cellular Biotechnology Training Program (T32GM145304). This work was initiated during the 2023 Quantitative Biology program at the Kavli Institute for Theoretical Physics (KITP), supported by grant NSF PHY-1748958 and the Gordon and Betty Moore Foundation Grant No. 2919.02.

## Data and Code Availability

All code used for analysis is available at github.com/idse/signaldecoding. The iterative IF data in this paper can be visualized online at: https://cdb-rshiny.med.umich.edu/2Dgastruloid_IIF/

## Methods

### Cell culture

Human pluripotent stem cells (ESI017; XX, ESI BIO) were cultured in mTeSR1 pluripotency maintenance medium (STEMCELL Technologies #85850) on cultrex-coated tissue culture plates. Medium was changed daily. Cells were passaged as whole colonies with L7^54^ when they reached 60-80% confluency, every 3-7 days. Cells were routinely checked for mycoplasma contamination and found to be negative.

### 2D gastruloid differentiation and drug treatment

Cells were disassociated using Accutase and resuspended as single cells. Cells were counted with a hemacytometer and seeded on micropatterned and laminin-coated substrates at 470k cells per square centimeter in mTeSR1 medium supplemented with 10 μM of the ROCK inhibitor Y-27632 (RI, MedChemExpress, HY-10583). After 45 minutes, wells were washed once with PBS to remove cells not bound to the laminin-coated patterns and medium was replaced with fresh mTeSR1 with RI. One hour after washing, the medium was changed to remove RI and add BMP4 (R&D, 314BP050CF) to induce differentiation. After 48 hours of differentiation, the differentiation medium was removed and the 2D gastruloids were fixed by incubation with freshly diluted 4% paraformaldehyde for 20 minutes at room temperature. For YAP activation, we compared several doses of TRULI (MedChemExpress, HY-138489) and found that 100uM induces strongly nuclear YAP across the colony but also severely reduces cell number, making it hard to separate indirect effects of density from direct effects of YAP activation. In contrast, 30uM TRULI robustly activated YAP without significant changes in cell number, therefore, we used 30uM in figure 5. Similarly, for ERK perturbation, we used the MEK inhibitor PD0325901 (SelleckChem, 50-136-0870) at a dose of 0.5uM, which was sufficient to significantly reduce ERK and severely affect fate without leading to large reductions in cell number that we saw at higher doses. For Wnt inhibition, we used the Wnt ligand secretion inhibitor IWP2 (ApexBio, A3512-10) at a dose of 5 µM.

### Iterative immunostaining

Iterative immunostaining and imaging was performed according to the protocol in Freeburne et al.^20^, adapted from Gut el al.^21^. Briefly, after fixation, colonies were subjected to a ‘pre-treatment’ in which they were incubated in blocking buffer (3% donkey normal serum, 0.1% Triton X-100, and Penicillin/Streptomycin diluted with PBS-/-) at room temperature for 2 hours, washed 3 times with elution buffer (EB; distilled water with L-glycine, urea, guanidine hydrochloride, Tris(2-carboxyethyl) phosphine hydrochloride at pH 2.5) with 10 minutes incubation in EB between each wash, and then washed 3 times with PBS. For the first round of staining, cells were incubated in blocking buffer for 30 minutes at room temperature. Then a solution of primary antibodies (Supplemental Table 1) diluted in blocking buffer was added and cells were incubated on an orbital shaker at 60 rotations per minute for 3 hours at room temperature. After primary antibody incubation, cells were washed 3 x 15 minutes each with PBST (PBS with 0.1% Tween 20), then incubated at room temperature for 1 hour on the orbital shaker with DAPI (Thermo Fisher Scientific, D1306) and secondary antibodies (Supplemental Table 2) diluted in blocking buffer, protected from light. The sample was washed twice more for 10 minutes each with PBST, and stored in PBS. PBS was replaced with imaging buffer (IB; distilled water with N-acetyl-l-cysteine and NaOH, adjusted to a pH of 7.4) in each well immediately before imaging that well. For subsequent rounds of immunostaining and imaging, antibodies from the previous round were eluted by washing the sample 3 times with EB for 10 minutes each, then washing 3 times with PBS. Staining for subsequent rounds was done following the same protocol as for the first round but without Triton X-100 in blocking buffer.

### Microscopy

Fluorescence imaging was performed with a Nikon/Yokogawa spinning disk confocal microscope with a 40x silicon oil objective using NIS Elements AR software. In the first round of imaging, representative colonies were chosen to be imaged. Each colony was captured with a z stack of 30-40 μm with 1 μm spacing between z slices, starting slightly below the focal plane of the bottom-most cells in the colony and ending slightly above the top-most cells. Xy resolution was 0.325 microns per pixel. Each colony was captured with a 4×6 montage of fields of view with 15% overlap that were stitched into a single large field of view. A flat field correction was applied to each panel of the image grids to compensate for uneven illumination, after which the panels were stitched together using custom MATLAB scripts available on our github repository (github.com/idse/signaldecoding).

### Segmentation

We segmented nuclei in each z-slice of the DAPI image stack from the first round of imaging for each colony using a combination of Cellpose and Ilastik^55–57^. Nuclear masks in each z slice were then linked to build 3D nuclear masks^18^. We used the Ilastik Object Classification Workflow to identify and discard junk and mitotic cells in the nuclear segmentation. We then created masks in which to read out cytoplasmic intensity for each nucleus as follows. We used the Ilastik Pixel Classification Workflow on the SMAD2/3 stain in the first round to segment each voxel as part of a cell or background. In each z slice we ran a watershed operation using segmented nuclei in that z slice as seeds. We then dilated each nuclear mask by a set number of pixels and excluded the original nuclear mask to create a ring about each nucleus. We intersected this ring with the watershed basin for the given nucleus so it would not include pixels belonging to a neighboring cell, and with the SMAD2/3 channel segmentation so the mask would only include pixels that are within cells. For a given nucleus, we combined these 2D masks in the different slices in which the nuclear mask was linked into a 3D cytoplasmic mask. Some colonies in some stains have small spots of brightly fluorescing junk that do not correspond to nuclei. To exclude this non-cell fluorescence from the quantification, we used Ilastik pixel classification to create a mask of pixels containing bright junk on the maximal intensity projection of colonies and stains in which this problem was observed. Intensity values for cells overlapping the junk segmentation in a given channel were dropped.

### Alignment between rounds

The image data in later rounds was aligned to the first round using a rigid translation and rotation based on their DAPI stains. Because any rotation was very small, these steps were performed sequentially for efficiency. The translation was determined based on phase correlation of the maximal intensity projection (MIP) of the DAPI stains. The second round DAPI MIP was then rotated about the center of the colony in 0.1 degree increments from -1.5 to +1.5 degrees to find the degree of rotation with maximum image cross-correlation. Once an optimal x-y shift and rotation were determined for a colony in a given round, those operations were applied to each z slice in each channel using the MATLAB interp2 function. There was also some slight variation in the starting z position used to capture the colony in each round, and we found that there was often a small amount of colony shrinking in the z direction between rounds of immunostaining. To measure this shift and shrinking in z we calculated pairwise cross-correlation in the DAPI channel between individual z slices in round 1 and round n to find the slice in round n with highest correlation to each slice in round 1. We then performed a linear regression on these pairs of (zround1, zround2) to find a shift (intercept) and scale (slope) to scale the second round image stack onto the frame of reference of the first-round stack. The MATLAB interp3 function was used to shift the z stack so that the bottom of the colony was aligned in both rounds and to scale the stack so the colonies have the same spatial extent in z. If the number of z slices in the shifted and scaled stack was smaller than the number in the first round of imaging, then additional slices of NaN values were appended to the top of the stack so the image stacks had the same dimensions. If the number of z slices in the second stack after alignment was greater than in the first round, then the aligned second-round stack was truncated to the size of the first. These operations result in pixel-level alignment between rounds of imaging so that nuclear masks segmented from the first round of staining can be applied to read out fluorescent intensities in each subsequent round.

### Correction for illumination bias

We observed small fluctuations in overall illumination between colonies in some rounds. To determine the illumination bias for a given colony in round n, the background-subtracted average intensity of each immunofluorescence stain in that round was divided by the average of that stain over all colonies in the same condition, and this value was averaged over all stains in that round. This provides a scaling factor for the colony relative to the average in the same condition, averaged over multiple stains. To adjust the overall illumination, pixel values over background were divided by this scaling factor when quantifying single-cell intensity values.

### Single-cell quantification

For each identified cell, the nuclear intensity was taken as the mean intensity in the segmented nuclear mask and the cytoplasmic intensity is the mean intensity in the segmented cytoplasmic mask around the nucleus. Voxels with NaN padding from the image alignment steps or which fall within the segmented junk mask for the channel being quantified are not included in the average. Colony intensity scaling was applied before quantification. Nuclear to cytoplasmic ratio values were taken for each channel as nuclear intensity minus background divided by cytoplasmic intensity minus background, where ‘background’ is the average intensity in the image outside of the colony, excluding NaN padding and junk.

### Local background estimation

Some stains appeared to exhibit a non-specific stain with intensity that depends on local cell density. The variation in background intensity readout in non-expressing cells can lead to artifactual gradients and correlations, particularly for markers like PRDM1 with a low percentage of expressing cells. To estimate the non-specific background for each nucleus, we took the bottom 5th percentile intensity value in a sphere of 7.5 um radius about the centroid of the nuclear mask for each cell. Because this density-dependent local background is presumed to arise from non-specific antibody binding, only voxels within cells are included. The local background was subtracted from the nuclear intensity for nuclear-localized proteins (transcription factors; this includes all used fate markers, pSMAD1/5/9, and LEF1, but not YAP, SMAD2/3, β-catenin, pERK, and pAKT).

### Spatial profiles and positional error analysis

The center of each colony was determined as the average of x-y centroid locations of nuclei in the colony weighted by nuclear area. Cells were then binned by distance from the center into 50 bins containing equal numbers of cells, which lead to the outer bin being about one cell diameter wide. The average distance from the center in the outer bin was taken as the true radius of the colony and the edge distance of a cell was defined as the true colony radius minus the cell’s distance from center.

### Discretization of fate markers and cell fate classification

To assign cell fates we first thresholded all the marker gene expression into positive or negative. Thresholds were picked manually based on inspection of histograms and scatter plots. We then assigned fates to the cells based on marker gene expression as follows:

1. PGCLC = TFAP2C+ & SOX17+
2. mesoderm = TBXT+ & TBX6+
3. amnion = ISL1+
4. PSLC = TBXT+ & (TBXT > NANOG)
5. pluripotent = NANOG+ & SOX2+ & (NANOG > TBXT)
6. ectoderm = NANOG-& (SOX2 > TBXT)

These conditions were applied in order, so cells were assigned to first fate for which they met the conditions (although the conditions should be largely mutually exclusive). Markers were normalized to threshold before comparison so NANOG+ means NANOG > 1. The condition TBXT > NANOG is to handle double positive cells in the continuum between pluripotency and PSLC. Finally, in the drug perturbation some cells appeared that were ISL1-TFAP2C+SOX17- and we assigned these amnion fate by adding a final rule amnion = (amnion | TFAP2C+).

### Spatial Fate Distribution Analysis

To evaluate whether the model correctly predicts the spatial organization of cell fates within colonies, we computed radial fate histograms for both measured and predicted fates. Cells were partitioned into N_bins_=20 equal-frequency radial bins based on their distance from the colony center. For each radial bin b, we computed the fate composition as the fraction of cells assigned to each fate m:

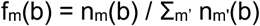

where n_m_(b) is the number of cells in bin b assigned to fate m. This yields a radial profile of fate proportions that can be directly compared between measured and predicted assignments.

### Fate Boundary Identification

To quantitatively compare the spatial extent of each fate, we identified radial boundary positions. For each fate m, the per-fate radial distribution was computed and normalized by its maximum value:

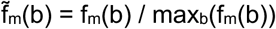

Boundary positions were defined as the radial locations where f̃_m_ crosses a threshold of 0.5. For each fate, we identified an inner boundary (where the normalized frequency rises above threshold from the colony edge) and an outer boundary (where it falls below threshold toward the colony center). Comparing measured and predicted boundary positions provides a quantitative assessment of how accurately the model captures the spatial organization of cell fates.

### Sig2Fate architecture and implementation

We employed a variational encoder-decoder network based on the Variational Information Bottleneck (VIB) framework^58,59^ to learn a compressed latent representation that maps single-cell signaling inputs to gene expression outputs. The model performs prediction at the single-cell level, taking each cell’s signaling state as input and predicting its gene expression profile. The architecture consists of three components: an encoder network, a stochastic latent bottleneck, and a decoder network.

The encoder transforms the signaling input vector into parameters of a latent distribution. Both encoder and decoder consist of D fully connected hidden layers with W neurons each and ReLU activation functions. We determined optimal architecture parameters W=32 and D=2 via cross-validation (Fig. S2a, b). Following the encoder, a bottleneck is introduced by two parallel linear output layers that produce the mean (μ) and log-variance (log σ²) of a 2-dimensional Gaussian distribution. Latent representations for each latent dimension i are sampled using

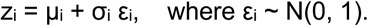

The stochasticity in the latent bottleneck injects noise during training to prevent overfitting and discontinuities in latent space. For model evaluation, predictions use only the posterior mean μ(s). Across independent training runs, we find similar distributions of points in latent space, up to translation and rotation. The decoder then maps the 2-dimensional latent representation to the 16-dimensional gene expression output space. The decoder architecture mirrors the encoder, consisting of D fully connected hidden layers with W neurons each and ReLU activations, followed by a linear output layer. The model was implemented in PyTorch.

### Sig2Fate training

The model is trained by minimizing the VIB objective^59^

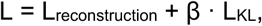

where the reconstruction loss is the mean squared error between predicted and observed gene expression levels for each cell:

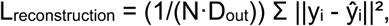

where D_out_=16 is the output dimensionality and the KL divergence regularizes the latent distribution toward a standard normal prior:

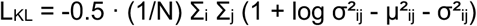

where i indexes over the N cells and j indexes over the latent dimensions. The KL divergence regularization explicitly penalizes deviation from the prior, forcing the encoder to transmit only information necessary for predicting gene expression while compressing out irrelevant signal variation. The hyperparameter β controls the trade-off between reconstruction fidelity and information compression in the latent space. We used β = 0.01. Both input signals and output gene expression values were z-score normalized (zero mean, unit variance) prior to training. Model parameters were optimized using the Adam optimizer with a learning rate of 10⁻³ for 800 epochs.

To assess model generalization and avoid overfitting, we employed k-fold cross-validation at the colony level (k=6 for the B50 condition). In each fold, one colony was held out as the test set while the remaining k-1 colonies were used for training. This colony-level splitting ensures that the model is evaluated on entirely unseen biological replicates rather than randomly sampled cells, providing a more stringent test of generalization.

### Evaluation of single cell gene predictions

To evaluate prediction accuracy at the gene level, each predicted gene expression value was thresholded into a binary classification (positive or negative) using the same thresholds applied to experimental data. This binary classification enables comparison between predicted and observed gene expression states for each cell, yielding four outcomes: true positives (TP), true negatives (TN), false positives (FP), and false negatives (FN). We computed the following metrics:

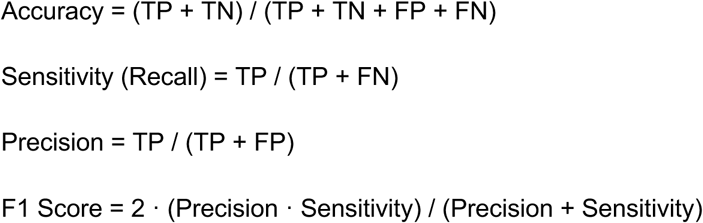

Accuracy measures the overall fraction of correct predictions. Sensitivity (also called recall) measures the fraction of true positives that were correctly identified. Precision measures the fraction of predicted positives that were correct. The F1 score is the harmonic mean of precision and sensitivity, providing a balanced measure of prediction performance.

For comparison, we derived the expected F1 score under random guessing. Consider a gene that is ON (positive) in a fraction p of cells. For informed random guessing, predictions are made with probability p of guessing positive (matching the true class frequencies). The expected outcomes are:

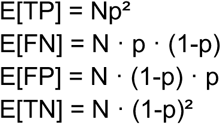

yielding:

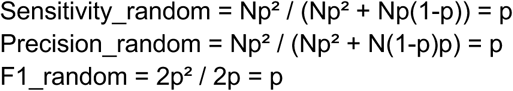

Thus, the expected F1 score under random guessing equals the fraction of positive cells p, providing a gene-specific baseline against which to compare model performance.

### Continuous gene prediction accuracy

To evaluate prediction accuracy for continuous gene expression values, we computed the conditional probability distribution P(g_exp_ | g_pred_), representing the distribution of experimentally observed gene expression g_exp given a predicted value g_pred. Predicted values were partitioned into N bins spanning the observed expression range. For each bin b of predicted values, we computed the conditional mean:

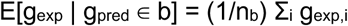

and conditional variance:

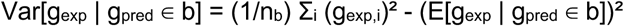

where the sum runs over the n_b_ cells whose predicted expression falls within bin b. Conditioning on predicted rather than observed values avoids regression dilution artifacts. For a perfect predictor, the conditional mean E[g_exp_ | g_pred_] equals g_pred_ (falling along the identity line) with zero variance.

To summarize prediction accuracy across all genes, we computed the gene-averaged conditional distribution:

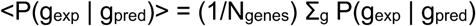

along with the gene-averaged conditional mean and variance. This provides an overall measure of how faithfully predicted expression values correspond to observed values across the full gene panel.

### Cell fate predictions

Cell fates were assigned from predicted gene expression profiles using the same thresholding procedure applied to experimental data (see Cell fate determination). This ensures consistency between experimentally observed and model-predicted fate assignments.

### Mutual information estimation

To quantify how much information the signaling state provides about each gene’s expression, we estimated the mutual information I(g; {s_j_}) between gene expression g and the set of N input signals {s_j_} = {s_1_, …, s_N_}. Direct estimation of mutual information in high-dimensional spaces is challenging due to the curse of dimensionality. We therefore used our regression predictions to obtain a lower bound via the Data Processing Inequality (DPI)^24^.

Consider the Markov chain g → {s_j_} → ĝ, where g is the observed gene expression, {s_j_} are the measured signals, and ĝ = f({s_j_}) is the predicted gene expression computed by the trained regression model. The DPI states that information can only be lost, never gained, through processing:

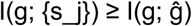

Thus, any decoder provides a lower bound on the true mutual information^27,60^. To derive the DPI, we apply the chain rule for mutual information to I(g; {s_j_}, ĝ) in two ways:

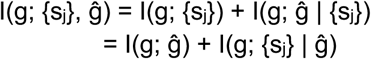

Since ĝ is a deterministic function of {s_j_}, knowing {s_j_} completely determines ĝ, so I(g; ĝ | {s_j_}) = 0. This gives:

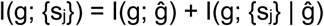

Since mutual information is non-negative, I(g; {s_j_} | ĝ) ≥ 0, yielding: I(g; {s_j_}) ≥ I(g; ĝ)

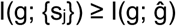

with equality if and only if ĝ is a sufficient statistic for g. The quantity I(g; ĝ) can be estimated directly from the joint distribution of observed and predicted gene expression values, providing a practical lower bound on the information that signals carry about each gene.

### Calculation of cumulative mutual information

Decoder-based MI between predicted and actual fate marker expression was estimated using scikit-learn’s mutual_info_regression, computed separately for each colony and averaged across colonies. Signaling features were added sequentially to build up a cumulative MI: at each round, the MI was computed for all combinations consisting of the previously selected features plus each remaining candidate feature. The candidate that gave the highest mean MI across colonies was selected, provided it was significantly better than all other candidates by paired t-test with Benjamini-Hochberg correction (α = 0.1); if no candidate was significantly better than all others, the candidate appearing earliest in the predefined signal ordering (pSMAD1, YAP, LEF1, β-catenin, SMAD2/3, SMAD2, pERK, pAKT) was selected as a tiebreaker. The cumulative MI at each round was then compared to the maximum MI achieved across all rounds by paired t-test; signals were considered to contribute significantly to the total if their inclusion brought the cumulative MI to a level not significantly different from the maximum (p > 0.05). This threshold is indicated by a dashed horizontal line in the cumulative MI plot, with error bars representing the standard error of the MI across colonies.

### Identification of an angle in signaling space

To relate polar coordinates in the latent space to polar coordinates in the signal space, we found two orthogonal vectors w1 and w2 in the standardized signal space that best approximate the latent coordinates L1 and L2 as linear combinations of the input signals. Here X denotes the matrix of z-scored input signals, with rows corresponding to cells and columns corresponding to the six signaling features (pSMAD1, YAP, LEF1, β-catenin, SMAD2/3, pERK). We minimized the mean squared error of predicting mean-centered L1 and L2 from mean-centered X, subject to the orthogonality constraint w1·w2 = 0, using sequential least-squares programming (SLSQP, scipy.optimize.minimize) with 20 random restarts. Signal-space analogs of the latent polar coordinates were then defined from the projections p1 = X·w1 + mean(L1) and p2 = X·w2 + mean(L2): a signal-space radius as the Euclidean distance from the centroid in the (p1, p2) plane, and a signal-space angle as φ_signal = arctan2(p2 - mean(p2), p1 - mean(p1)). Correspondence with the latent radius and angle φ were quantified by the Pearson and circular correlation coefficients respectively.

### Predicting perturbed signals with naive and k-nearest neighbor approaches

Both naive and k-nearest neighbor (knn) predictions are based on the same B50 data that was used for Figures 2-4 and use the same thresholds to estimate perturbed signaling: the 5th percentile value in the B50 population for ERK inhibition, 0.1 percentile for Wnt inhibition, and the 95th percentile for YAP activation. For Wnt inhibition we used a high drug dose to shut down all Wnt signaling, justifying setting LEF1 close to the minimum observed in control (while trying to exclude the lowest outliers by not taking the absolute minimum). In contrast, for ERK and YAP we used drug doses that did not maximally activate or inhibit the pathways to maintain cell viability so we chose a less extreme cutoff. The 5th percentile values correspond approximately to the signaling levels for these pathways in the amnion, which as a population has the lowest ERK and highest nuclear YAP levels in control while maintaining viability. For naive predictions, the signal being perturbed is set to a fixed value for all cells. All other signals are held fixed, and the B50-trained model is evaluated on the adjusted signals. For knn-predictions, we first smoothed the B50 data using a Gaussian weighted average of the 10 closest neighbors for each cell in signaling space to avoid projecting onto outlier cells. Next, we applied the same percentile thresholds as above to identify a subset of B50 cells hypothesized to reflect the perturbed signaling state. We then projected each cell onto this subset by computing the z-normalized signaling distance between cells (excluding the perturbed signal) and replacing each cell with the Gaussian-weighted average of its 5 nearest neighbors in the subset. For ERK and Wnt inhibition, since these signaling pathways are known to be downstream of BMP, we exclude pSMAD1 in this process of knn-adjustment and keep it unchanged from B50.

### AI use

Anthropic’s Claude was used as an aid in preparing this manuscript: for editing and stylistic revision of the text, and for assistance in writing portions of the analysis code. All code was reviewed and tested by the authors. All scientific content, analyses, and conclusions are the authors’ own.

## Supplemental Tables

**Table 1:** Primary Antibodies.

| Antigen | Manufacturer | Catalog number | Dilution |
| --- | --- | --- | --- |
| β-catenin | BD Biosciences | BDB610154 | 1/200 |
| CDX2 | BioGenex | MU392A-5UC | 1/50 |
| EOMES | R&D Systems | MAB6166 | 1/200 |
| FOXA2 | R&D Systems | AF2400 | 1/500 |
| GATA3 | SantaCruz Biotechnology | sc-268 | 1/50 |
| HAND1 | R&D Systems | AF3168 | 1/200 |
| ISL1 | DSHB | 39.4D5 | 1/200 |
| LEF1 | Cell Signaling Tech | C12A5 | 1/100 |
| MIXL1 | ABclonal | A17223 | 1/150 |
| NANOG | R&D Systems | AF1997 | 1/100 |
| OCT3/4 | BD Biosciences | 611202 | 1/400 |
| OTX2 | R&D Systems | AF1979 | 1/150 |
| pAKT (Ser473) | Cell Signaling Tech | 4060 | 1/200 |
| pERK | Cell Signaling Tech | 9101S | 1/100 |
| PRDM1 | SantaCruz Biotechnology | sc-47732 | 1/25 |
| SMAD1 | Cell Signaling Tech | 6944T | 1/100 |
| SMAD2 | Cell Signaling Tech | 5339T | 1/50 |
| pSMAD1/5/9 | Cell Signaling Tech | 13820S | 1/50 |
| SMAD2/3 | BD Biosciences | 610843 | 1/50 |
| SNAI1 | R&D Systems | AF3639 | 1/100 |
| SOX17 | R&D Systems | AF1924 | 1/1000 |
| SOX2 | Cell Signaling Tech | 3579S | 1/200 |
| TBX6 | R&D Systems | AF4744 | 1/100 |
| TBXT | R&D Systems | AF2085 | 1/300 |
| TFAP2C | SantaCruz Biotechnology | sc-12762 | 1/150 |
| YAP | Cell Signaling Tech | 14074T | 1/100 |

**Table 2:**
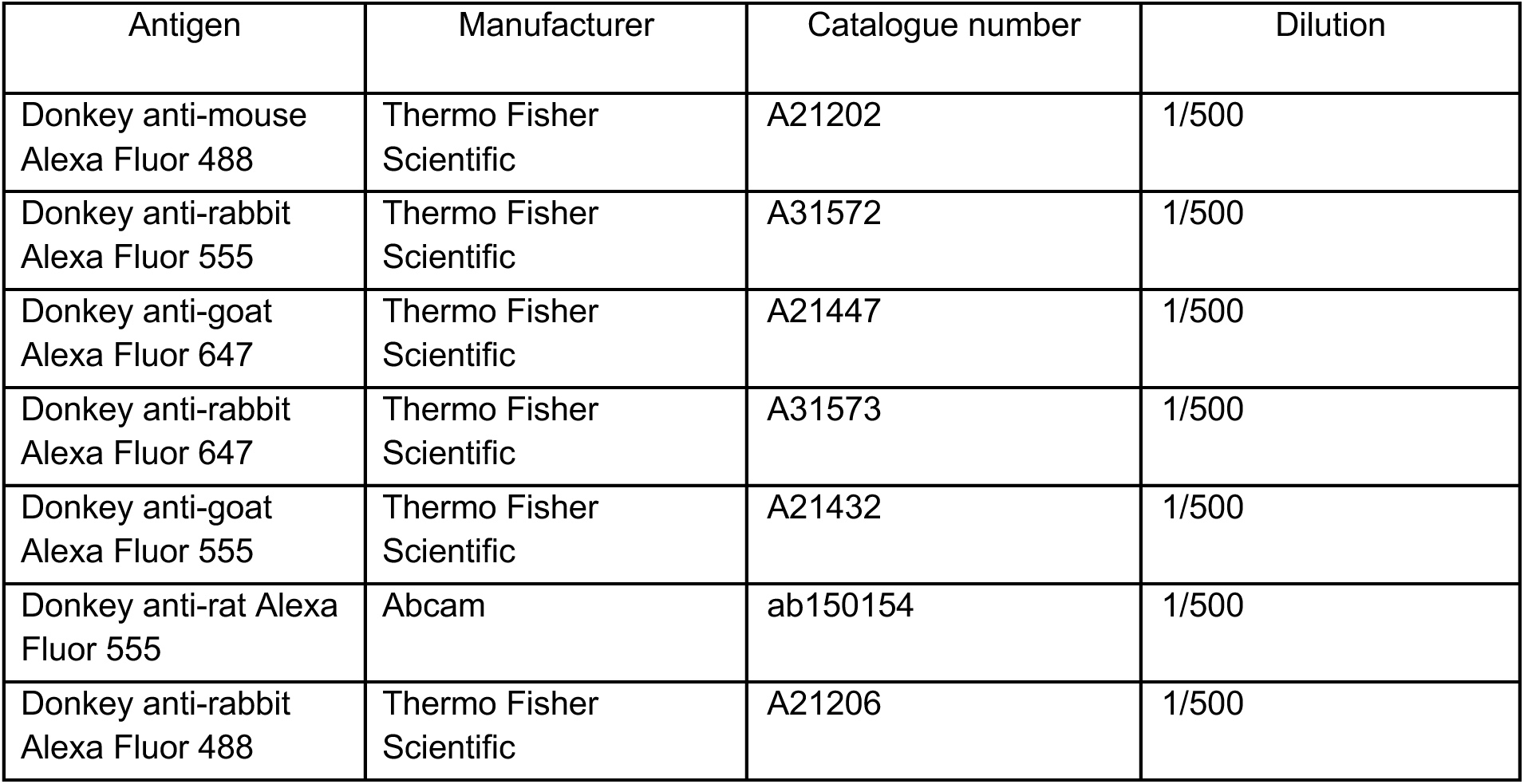
Secondary Antibodies.

| Antigen | Manufacturer | Catalogue number | Dilution |
| --- | --- | --- | --- |
| Donkey anti-mouse<br>Alexa Fluor 488 | Thermo Fisher<br>Scientific | A21202 | 1/500 |
| Donkey anti-rabbit<br>Alexa Fluor 555 | Thermo Fisher<br>Scientific | A31572 | 1/500 |
| Donkey anti-goat<br>Alexa Fluor 647 | Thermo Fisher<br>Scientific | A21447 | 1/500 |
| Donkey anti-rabbit<br>Alexa Fluor 647 | Thermo Fisher<br>Scientific | A31573 | 1/500 |
| Donkey anti-goat<br>Alexa Fluor 555 | Thermo Fisher<br>Scientific | A21432 | 1/500 |
| Donkey anti-rat Alexa<br>Fluor 555 | Abcam | ab150154 | 1/500 |
| Donkey anti-rabbit<br>Alexa Fluor 488 | Thermo Fisher<br>Scientific | A21206 | 1/500 |

**Table 3:** Staining order for experiments in figures 2-4 and corresponding supplements.

| Staining Round | Secondary antibody<br>dye: Alexa 488 | Secondary antibody dye:<br>Alexa 555 | Secondary antibody<br>dye: Alexa 647 |
| --- | --- | --- | --- |
| 1 | SMAD2-3 (mouse) | pAKT (rabbit) | SOX17 (goat) |
| 2 | GATA3 (mouse) | OTX2 (goat) | LEF1 (rabbit) |
| 3 | ISL1 (mouse) | TBXT (goat) | pSMAD1/5/9 (rabbit) |
| 4 | EOMES (mouse) | YAP (rabbit) | FOXA2 (goat) |
| 5 | TFAP2C (mouse) | NANOG (goat) | pERK (rabbit) |
| 6 | BCAT (mouse) | TBX6 (goat) | SMAD2 (rabbit) |
| 7 | SOX2 (rabbit) | PRDM1 (rat) | SOX17 (goat) |
| 8 | OCT3/4(mouse) | SMAD1 (rabbit) | HAND1 (goat) |
| 9 | CDX2 (mouse) | MIXL1 (rabbit) | SNAI1 (goat) |

**Table 4:** Staining order for LATSi experiment in figure 5 and corresponding supplements.

| Staining Round | Secondary antibody<br>dye: Alexa 488 | Secondary antibody dye:<br>Alexa 555 | Secondary antibody<br>dye: Alexa 647 |
| --- | --- | --- | --- |
| 1 | ISL1 (mouse) | TBXT (goat) | pAKT (rabbit) |
| 2 | SMAD2-3 (mouse) | SOX17 (goat), used | LEF1 (rabbit) |
| 3 | BCAT (mouse) | NANOG (goat) | pSMAD1/5/9 (rabbit) |
| 4 | TFAP2C (mouse) | TBX6 (goat) | pERK (rabbit) |
| 5 |  | SOX17 (goat) | YAP (rabbit) |
| 6 |  | SOX17 (goat) | SOX2 (rabbit) |

**Table 5:** Staining order for MEKi and WntSeci experiments in **figure 5** and corresponding supplements.

| Staining Round | Secondary antibody<br>dye: Alexa 488 | Secondary antibody dye:<br>Alexa 555 | Secondary antibody<br>dye: Alexa 647 |
| --- | --- | --- | --- |
| 1 | ISL1 (mouse) | TBXT (goat) | pERK (rabbit) |
| 2 | SMAD2-3 (mouse) | SOX17 (goat), used | LEF1 (rabbit) |
| 3 | BCAT (mouse) | NANOG (goat) | pSMAD1/5/9 (rabbit) |
| 4 | TFAP2C (mouse) | TBX6 (goat) | pAKT (rabbit) |
| 5 |  | SOX17 (goat) | YAP (rabbit) |
| 6 |  | TBX6 (goat), used | SOX2 (rabbit) |
| 7 |  |  | SOX2 (rabbit), used |

## Supplemental Figures

**Fig. S1:**
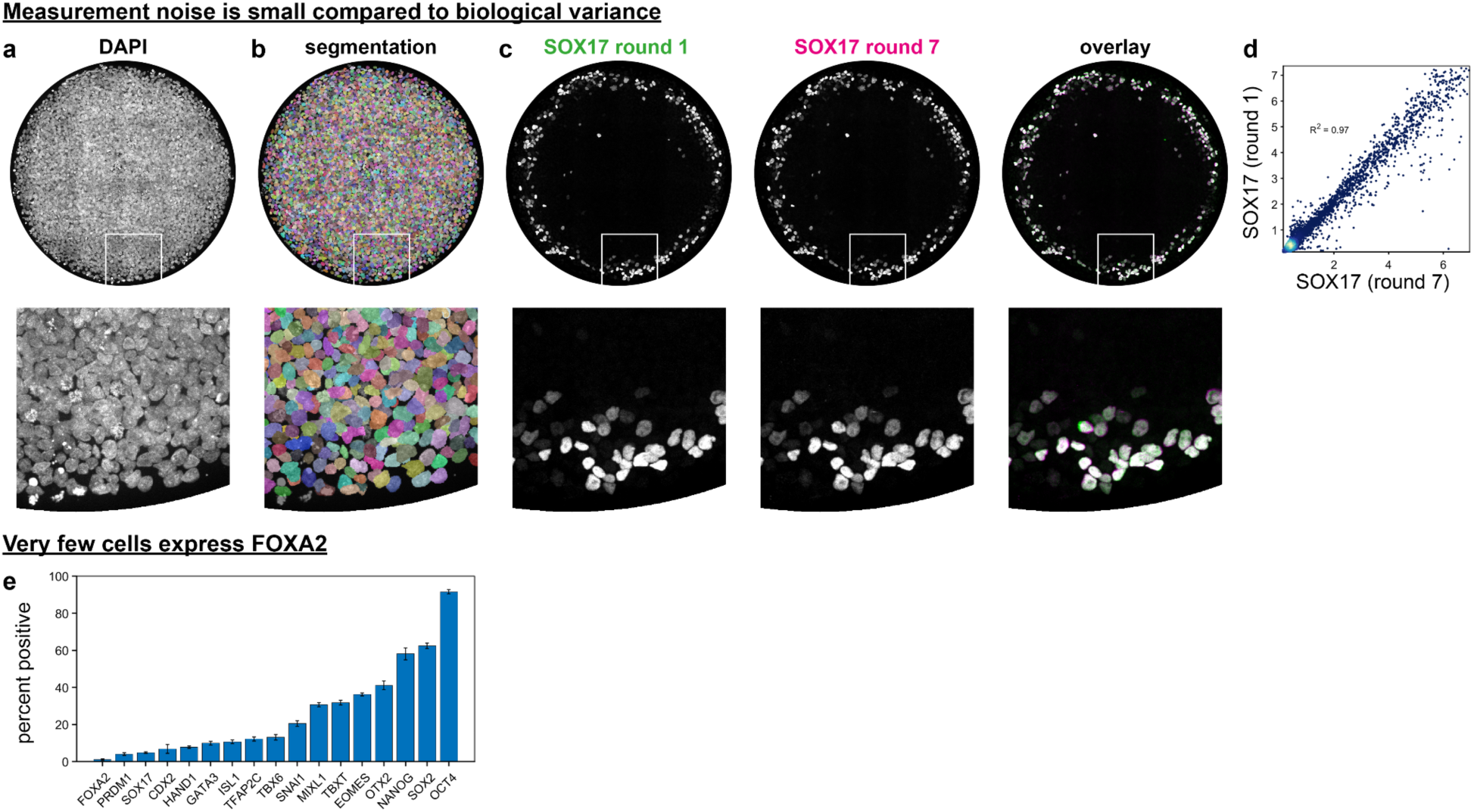
Image processing and characterizing technical vs biological noise. **a)** DAPI stain of example colony. **b)** Overlay of segmentation on DAPI. **c)** Round 1 and 7 SOX17 stains and overlay. White box in a-c is shown magnified below. **d)** Scatterplot of single cell SOX17 intensities in round 1 versus round 7 with fraction of variance explained R^2^=0.97. **e)** Fraction of cells expressing each of the marker genes (see methods).

**Fig. S2:**
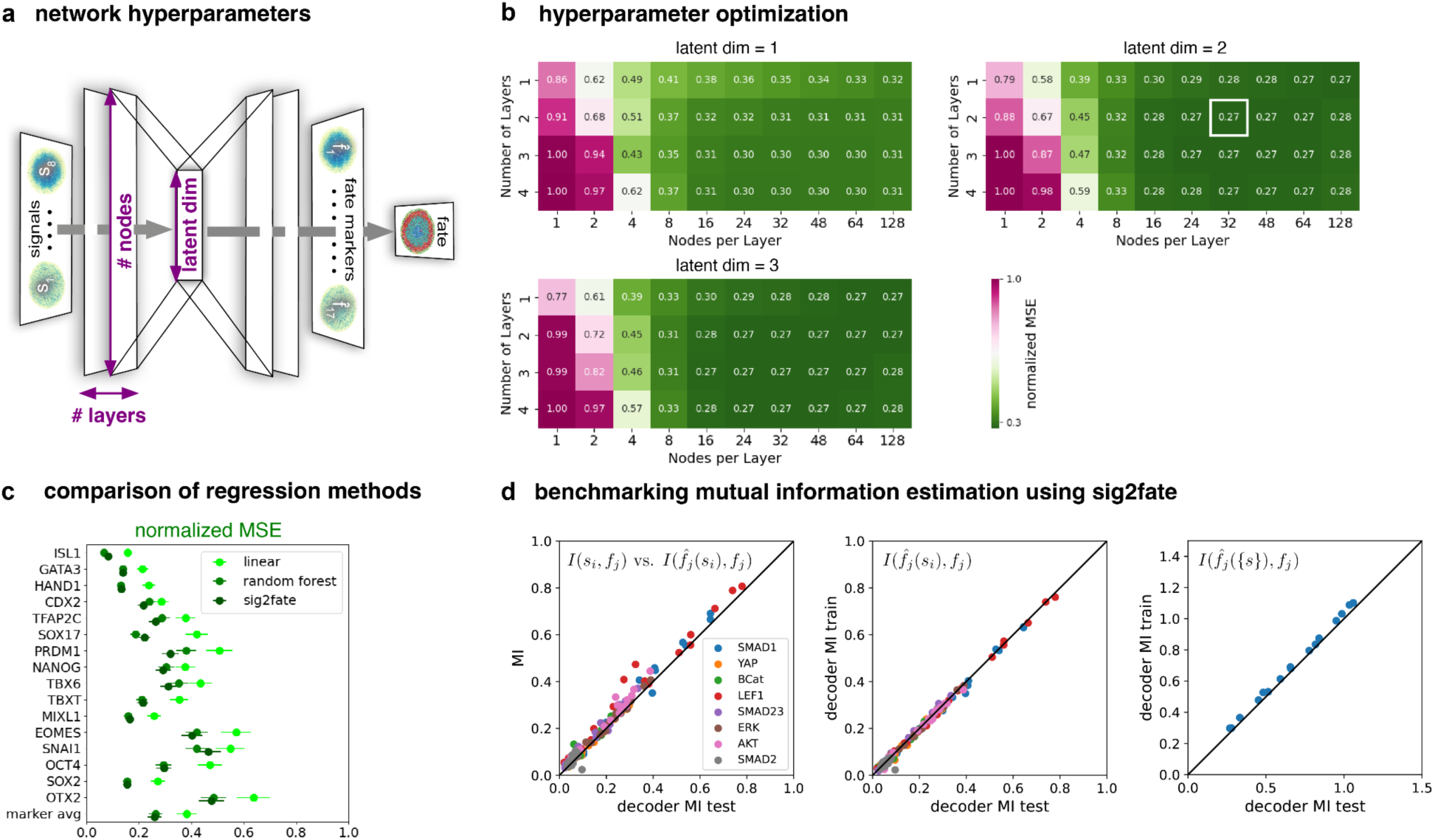
Optimizing hyperparameters, regression algorithms and benchmarking mutual information estimation (B50). **a)** Schematic of the NN architecture with hidden layers, nodes per layer, and bottleneck (latent) dimension as hyperparameters. **b)** Sweeps over numbers of layers and nodes per layer as well as latent dimension evaluating the mean squared error (MSE) for marker genes (after z-scoring and normalized to maximal error). White box indicates the hyperparameter values used throughout the paper. **c)** MSE for Sig2Fate with optimized hyperparameters compared with linear and random forest regression. **d)** Mutual information between signals and genes. Left: Decoder-based mutual information between single signals and genes by training Sig2Fate to predict all fate markers from individual signals is compared to direct calculation of mutual information using sklearn’s standard k-nearest-neighbor estimator. Middle: decoder-based mutual information between single signals and genes in test and training data sets. Agreement indicates that the model did not overfit to the training data. Right: mutual information between all signals combined and individual genes in test and training data sets.

**SI Fig. S3:**
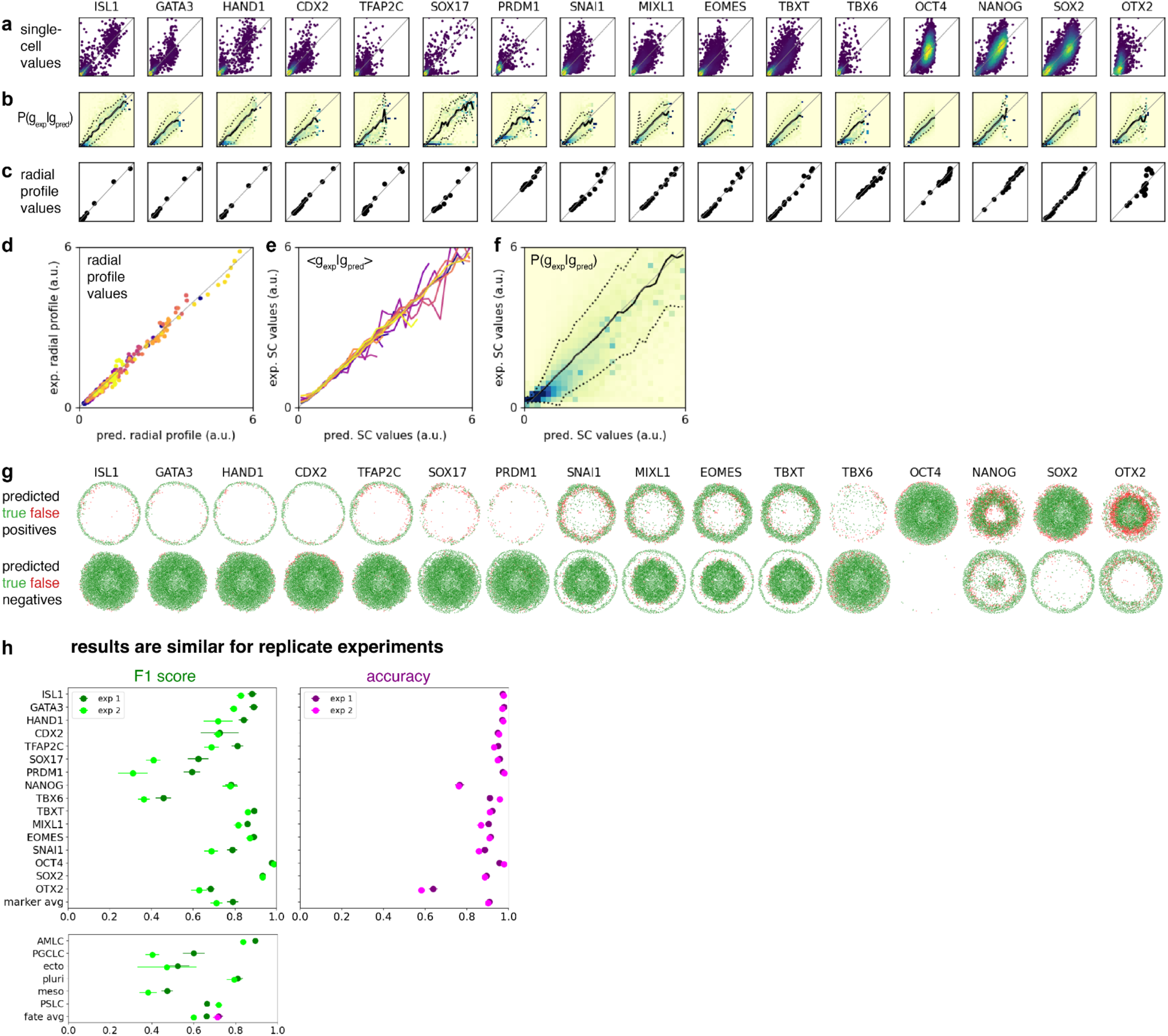
Evaluating model performance (B50). **a)** Single-cell gene values, experiment (y-axis) vs model prediction (x-axis). **b)** Conditional probability distribution P(g_exp|g_pred). Solid lines: conditional mean <g_exp|g_pred>. Dashed lines: standard deviation. **c)** Radial profile values, experiment (y-axis) vs model prediction (x-axis). **d)** All radial profile values plotted together. **e)** All conditional means plotted together. **f)** Conditional probability distribution P(g_exp|g_pred) averaged over all genes. Solid lines: conditional mean <g_exp|g_pred>. Dashed lines: standard deviation. **g)** True and false positives and negatives for thresholded prediction of gene expression. From these, the accuracy and F1 score in Fig. 2 are calculated. **h)** F1 score and accuracy for two replicate experiments.

**SI Fig. S4:**
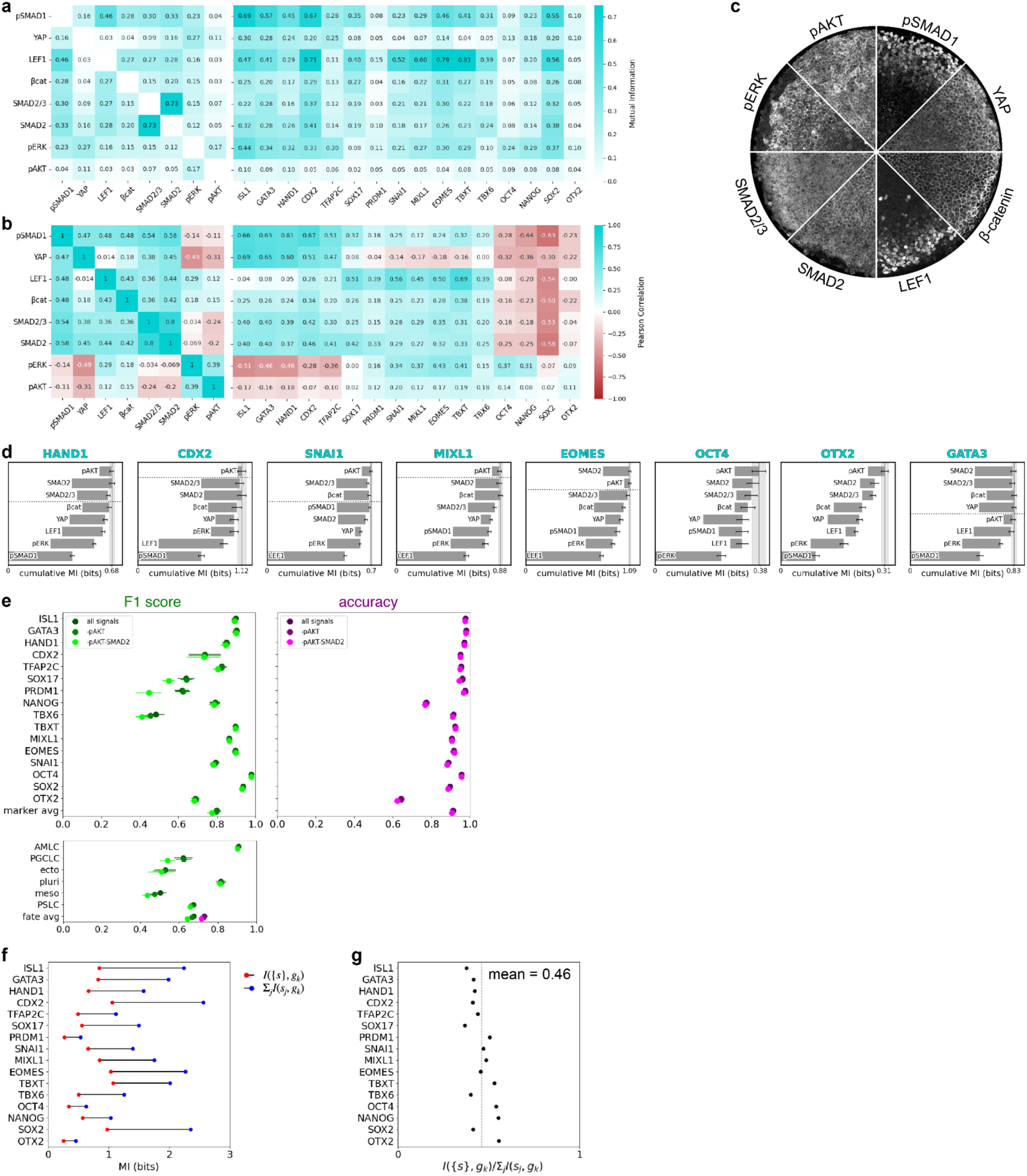
Quantifying combinatorial information in signal to gene mapping. **a)** Pairwise mutual information for signal-signal (left) and signal-gene (right) mutual information. **b)** Signal-signal (left) and signal-gene (right) correlations. **c)** Immunofluorescence stains for all 8 signaling readouts shown as slices of the same colony, same as in Fig.3d. **d)** Cumulative mutual information with an increasing number of signals (along the y-axis) for all genes. Length of each bar indicates the mutual information of each single signal with the gene. Color code corresponds to that used in panel b. **e)** Model performance after removal of pAKT, β-catenin, and SMAD2. **f)** Comparison of combinatorial mutual information of all signals with each gene (red dots) against the sum of all single-signal mutual information values with that gene (blue dots) in the reduced model. **g)** Combinatorial mutual information divided by summed single-signal mutual information values. The mean across genes is indicated by a dashed line.

**SI Fig. S5:**
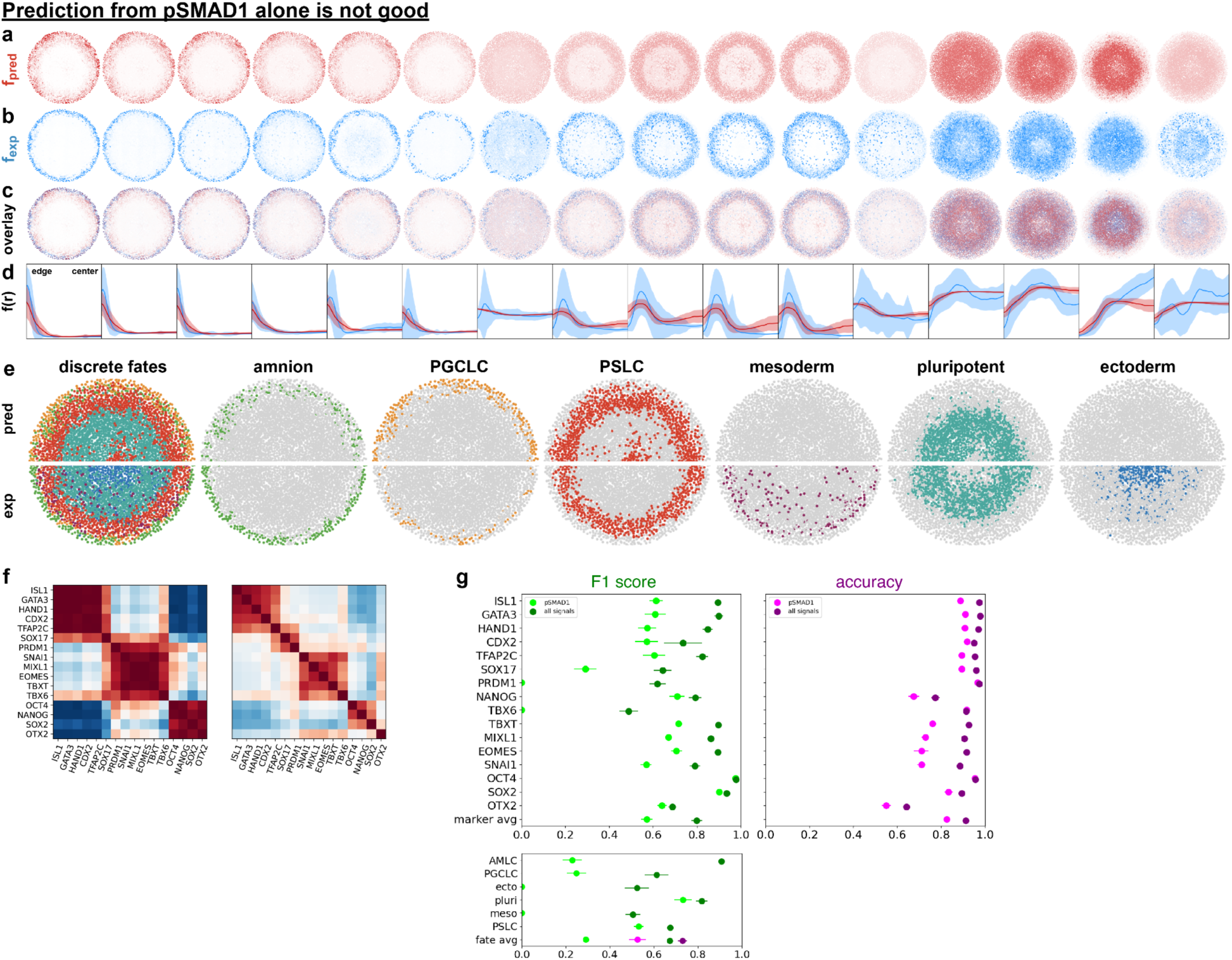
Prediction from pSmad1 alone. **a)** Fate marker genes predicted by a model trained from pSmad1 alone. **b)** Measured fate marker gene expression. **d)** Predicted (red) and experimental (blue) radial profiles. Shaded areas indicate standard deviation around the mean. **e)** Predicted (top half) vs measured (bottom half) discretized cell fate patterns. **f)** Predicted (left) and measured (right) gene-to-gene correlation matrix. **g)** F1 score and accuracy for pSMAD1 only model versus full model with all measured signals.

**SI Fig. S6:**
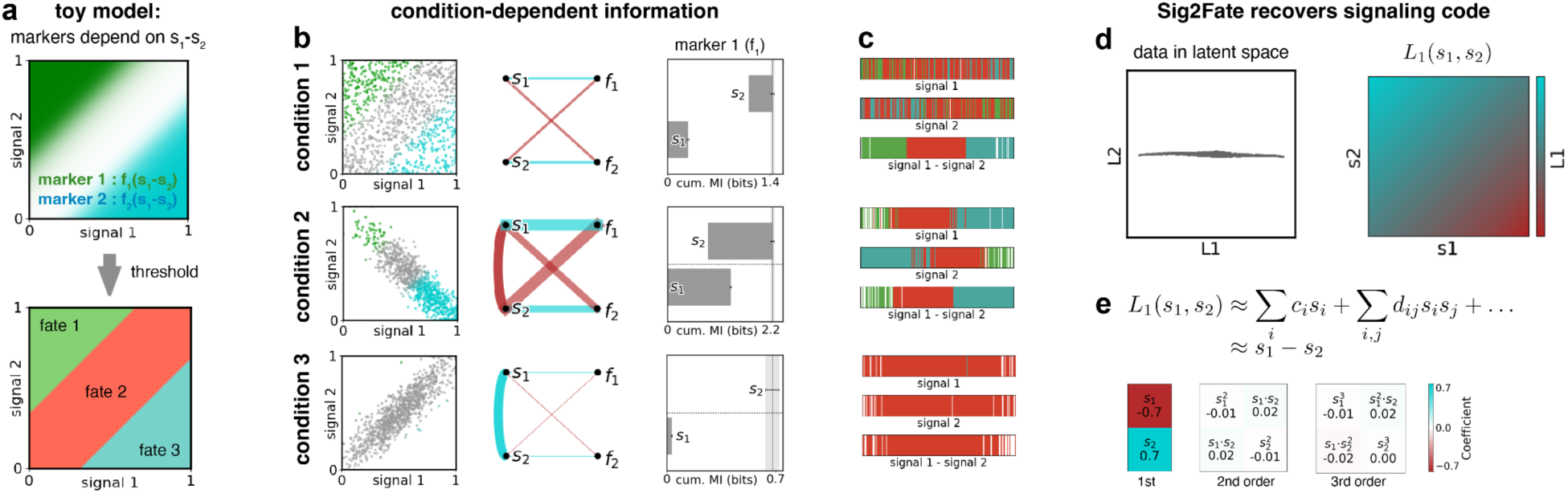
Toymodel shows how Sig2Fate recovers the relationship between signals and fate. **a)** In our toy model, expression of fate marker genes depends only on the difference between two signals. **b)** The pair-wise signal-signal and signal-fate mutual information (middle), as well as the mutual information between the combined signals and fate (right), depend strongly on how the data is distributed in the signaling space, even though the underlying relationship between signals and fate is fixed. **c)** This can be visualized intuitively by plotting how well cells separate by fate when sampled randomly in equally spaced bins along each signaling axis. 1000 cells per condition, 200 bins, white bins are empty. **d)** However, regardless of distribution, data from all three conditions ends up on a line through the Sig2Fate latent space, indicating a one-dimensional latent space (left), and the latent coordinate along this line (L1), appears to vary as a function of the signaling difference. **e)** A polynomial fit up to third order confirms that the latent coordinate L1 is proportional to the signal difference: Sig2Fate has correctly recovered the underlying relationship between signals and fate.

**SI Fig. S7:**
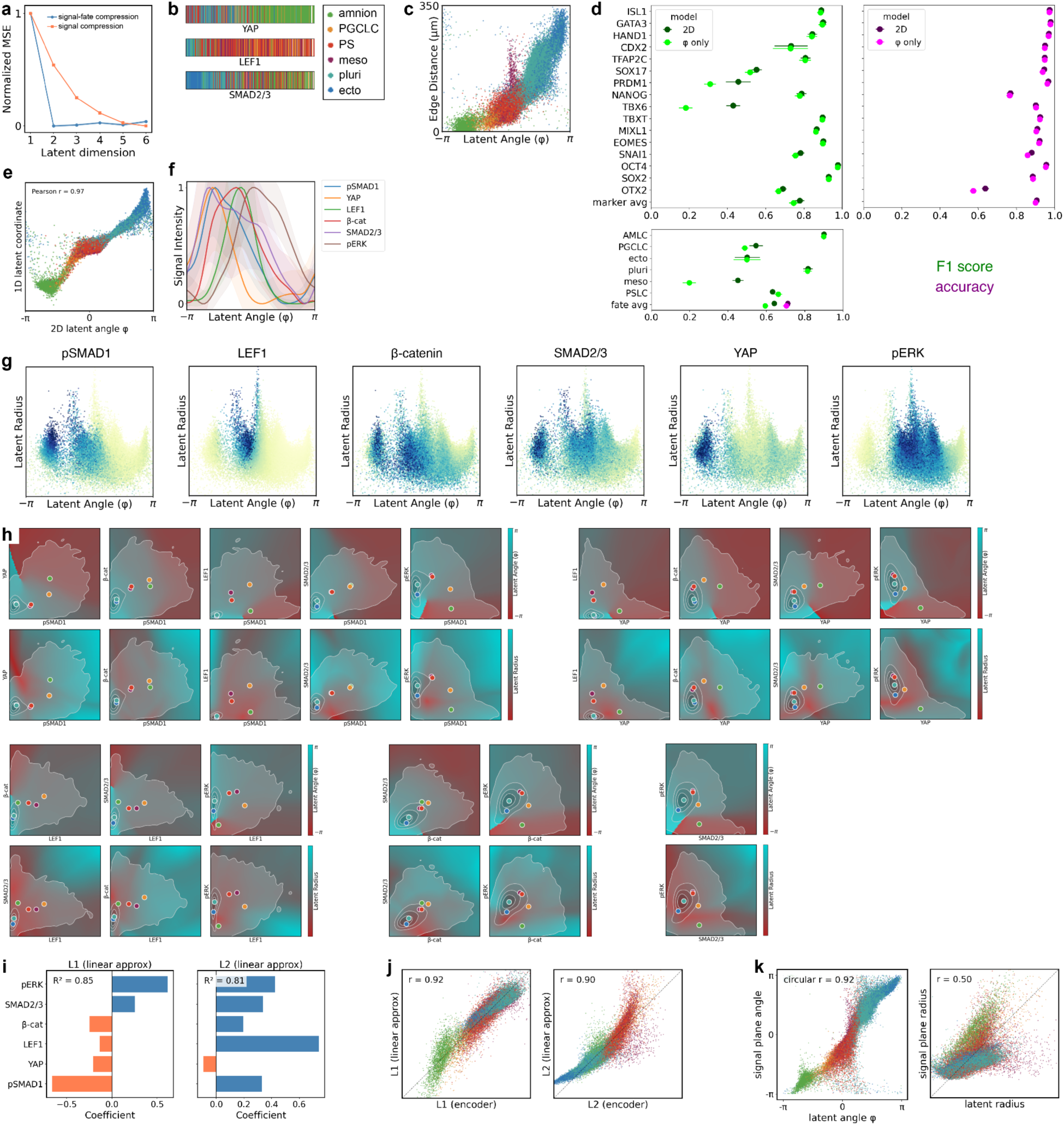
Interpreting the signaling code. **a)** Normalized mean squared error (MSE) for signal-to-fate and signal-to-signal encoder-decoders as a function of latent dimension with architecture otherwise as in other figures (2 hidden layers of 32 nodes). **b)** Bar plot showing fate for randomly sampled cells in equally spaced bins across the range of each signal, N=450 cells total, upper and lower limits are 1st and 99th percentiles. **c)** Scatter plot of latent angle versus edge distance colored for cell fate. Fate colors as in (b). **d)** Prediction performance of full model versus latent angle only, where in the latter radius is set to the average of all cells for that angle. **e)** Latent coordinate on 1D latent space versus angle of the 2D latent space have a correlation coefficient of 0.97. **f)** Mean signaling as a function of latent angle, error bands represent standard deviation. **g)** Scatter plot of data in latent polar coordinates colored for different signals. Color limits: 1st and 99th percentiles. **h)** Heatmap of latent angle and radius for all pairs of input signals, evaluated at the mean for all other signals conditional on the pair of signals being plotted, overlaid with kernel density estimate of data density (contour lines) and the mean of each fate (colored dots, colors as in (a)). **i)** Coefficients for linear regression of 2D latent coordinate L1 and L2 against the input signals. **j)** Scatter plots of linear approximation of latent coordinates against true latent coordinates. **k)** Angle and radius in the plane in signaling space defined by the linear approximation of the 2D latent coordinates plotted against the angle and radius on the latent space.

**SI Fig. S8:**
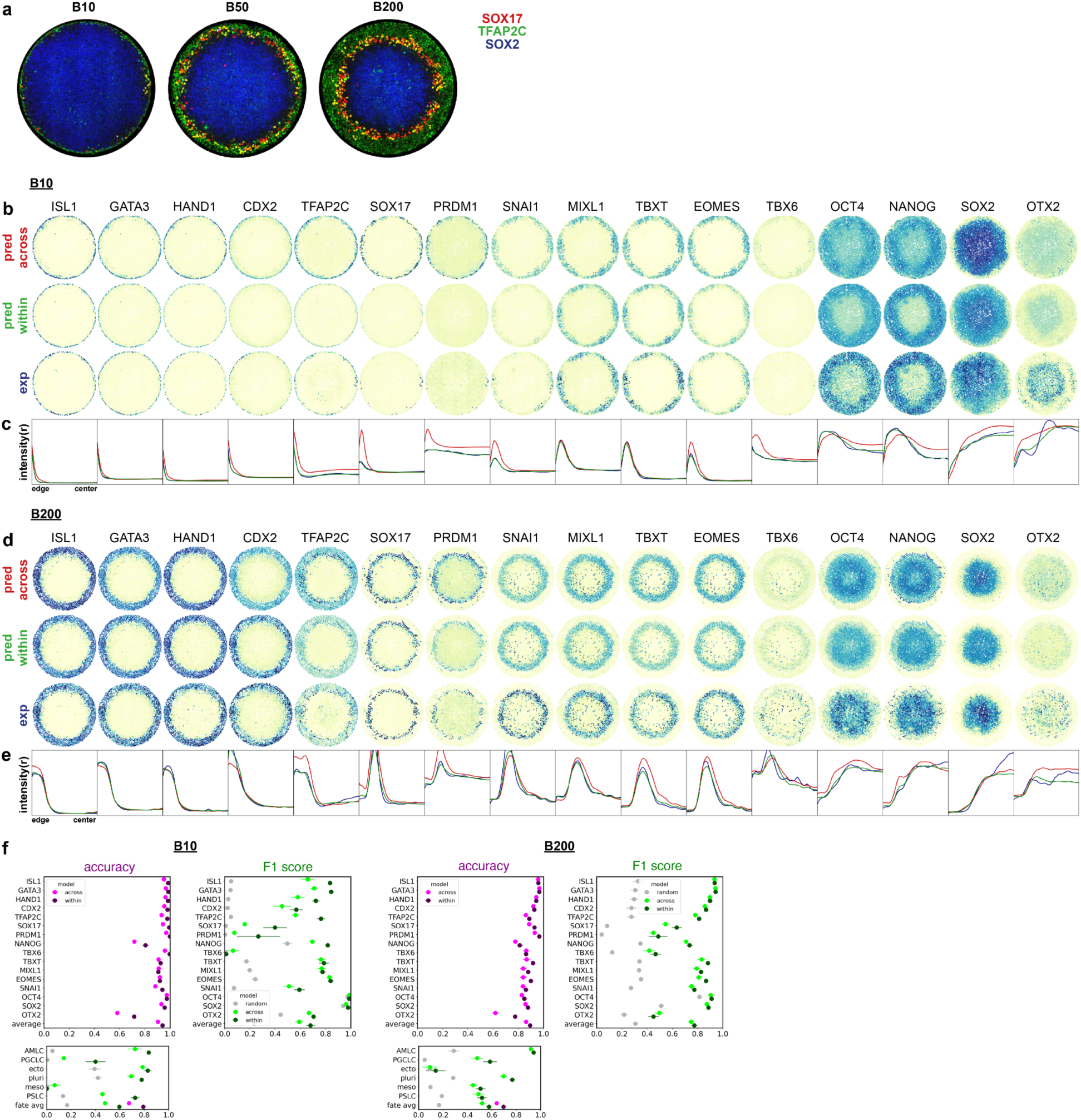
Fate prediction within vs. across conditions. **a)** IF data showing the distribution of PGCLCs (SOX17+TFAP2C+) in the same colonies as shown in Fig.4de. **b)** Fate marker genes predicted for B10from a model learned on B50 and evaluated across conditions (top); from a model learned within each condition (middle); and measured (bottom). **c)** Radial profiles predicted across conditions (red), within conditions (green) and measured (blue). **d,e)** Like b,c but for B200. **f)** Single cells prediction performance within and across conditions.

**SI Fig. S9:**
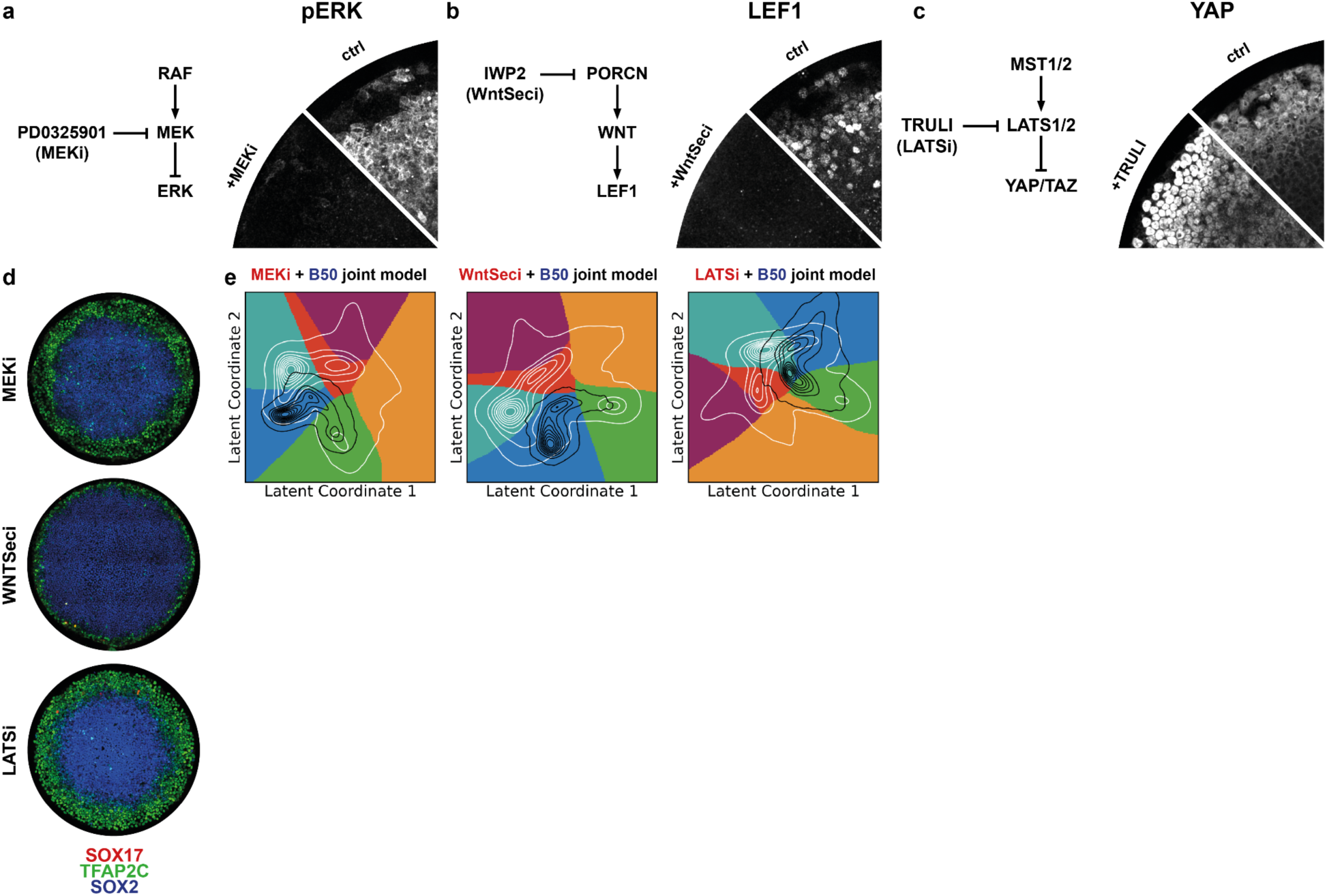
Predicting drug perturbations. **a-c)** Left: Simplified diagrams relating the drug treatments to measured signaling readouts. Right: stain for the relevant signaling readout after treatment vs. control. **d)** IIF stains of fate marker expression patterns for the same colonies as in (Fig.5c). **e)** Latent space visualization of the fate map for joint models trained on both control (B50) and perturbation data, overlaid are contours for the density of cell in control (white) and perturbation (black).

